# BASP1 Couples Ca^2+^ Signaling and Actin Polymerization to Mitochondrial Fission Essential for Neurite Outgrowth

**DOI:** 10.1101/2025.09.05.674494

**Authors:** Ekaterina Grebenik, Sofia Zaichick, Angel Gomez, Gabriela Caraveo

## Abstract

Actin-mediated mitochondrial fission is essential for cellular homeostasis, yet the mechanisms by which actin is recruited to mitochondria and how it couples the outer and inner mitochondrial membranes (IMM) remain poorly understood. Using a phosphoproteomic screen in a rat model of α-synucleinopathy, we identified BASP1 as a calcineurin-dependent substrate that is constitutively dephosphorylated under pathological Ca^2+^ elevations and phosphorylated under neuroprotective calcineurin inhibition. Immunoprecipitation and mass spectrometry of phosphomutant BASP1 expressed in neurons revealed that dephosphorylation promotes interactions with actin and IMM proteins. Dephosphorylated BASP1 recruits actin to mitochondria, while subsequent phosphorylation enables actin-mediated mitochondrial fission and neurite elongation. Constitutive dephosphorylation, as it occurs in α-synucleinopathy, impairs mitochondrial fission, inhibits neurite growth and promotes α-synuclein aggregation. Our findings position BASP1 as a Ca^2+^-CaN-regulated hub that coordinates actin remodeling and couples mitochondrial membranes to drive fission, revealing a mechanistic axis linking mitochondrial dysfunction to neuronal morphogenesis and α-synuclein pathobiology.

## INTRODUCTION

Mitochondrial fission and fusion are central to mitochondrial dynamics, regulating the organelle shape, number, distribution, and function (*1*). These processes are essential for mitochondrial quality control, subcellular localization, and cellular energy homeostasis. Mitochondrial fission is defined by two main events: an initial constriction of mitochondrial membranes and the subsequent fragmentation. In mammalian cells, fragmentation is mediated by the dynamin-related protein (Drp1) which self-assembles on the surface of the outer mitochondrial membrane (OMM) into oligomeric rings which constrict and divide the organelle (*2–4*). Prior to Drp1 recruitment, the initial constriction step, which is necessary for Drp1 assembly, occurs at ER-mitochondria contacts where ER tubules associate wrap around the OMM (*5–7*). Along these contact zones, actin filaments polymerize, providing the force needed for constriction and Drp1 recruitment (*2, 6, 8–17*). Although actin polymerization is critical for the first constriction step and known to be triggered by elevations in Ca^2+^ (*13*), the precise mechanisms by which actin is targeted to mitochondria, and how it coordinates fission between the OMM and inner mitochondrial membrane (IMM), remain poorly understood.

Disruptions in mitochondrial dynamics have been implicated in numerous neurological disorders, including Alzheimer’s disease, Parkinson’s disease (PD), Huntington’s disease, hereditary ataxias, and Charcot-Marie-Tooth disease (*12, 18–20*). A hallmark of both sporadic and familial forms of PD is the misfolding of α-synuclein (α-syn) (*21, 22*). Our group and others have previously identified a central role for the Ca^2+^-dependent phosphatase calcineurin (CaN) in α-syn pathobiology (*23–29*). We demonstrated that disease-associated forms of α-syn induce a pathological increase in cytosolic Ca^2+^, leading to aberrant activation of CaN ultimately triggering neuronal death (*23, 24*). However, partial inhibition of CaN with sub-saturating doses of FK506, a specific CaN inhibitor, rescues neurons from α-syn-induced toxicity (*23, 24, 28*). To uncover CaN-dependent substrates involved in α-syn pathobiology, we performed a phospho-proteomic screen in an *in vivo* rat model of α-synucleinopathy treated with either FK506 or vehicle (*23*). From this analysis, brain acid-soluble protein 1 (BASP1), also known as NAP-22 or CAP-23, emerged as one of only two proteins post-translationally regulated by CaN in response to FK506 *in vivo* (*23*).

BASP1 is a ubiquitously expressed protein upregulated during periods of axonal growth and synaptogenesis, where it plays a critical role in neurodevelopment (*30–32*). Homozygous knockout of BASP1 results in high neonatal lethality, with only 5–10% of animals surviving to adulthood (*33*). Surviving adults display significant neurodevelopmental impairments, including enlarged brain ventricles, axonal and synaptic abnormalities in the neocortex and hippocampus, and hyperactive behavior. Neurons lacking BASP1 display impaired neurite outgrowth and defects in the actin cytoskeleton (*33*). Although BASP1 is downregulated in most brain regions following the completion of axonal innervation, it remains highly expressed in the neocortex and hippocampus, suggesting a role in learning and memory (*34*). Notably, BASP1 expression is dynamically regulated in adulthood, especially during nerve regeneration after injury (*31, 33, 35*). Although BASP1 is known to regulate the actin cytoskeleton, the mechanisms by which it modulates actin dynamics in the adult brain remain unclear. Further, a recent study have linked BASP1 to PD. A meta-analysis of postmortem brain data from idiopathic PD patients identified BASP1 as a key regulator of PD-related genes (*36*). Here, using an unbiased phosphoproteomic screen in a rat model of α-syn pathology, we found that serines 40 and 165 (corresponding to human S40 and S170, respectively) of BASP1 are dephosphorylated under α-syn-induced pathological conditions and phosphorylated following neuroprotective treatment with sub-saturating doses of the CaN-inhibitor, FK506. Analysis of S40 and S170 phosphomutants of BASP1 in primary embryonic rat cortical neurons revealed that phosphorylation at these sites regulates BASP1 interactions with α-syn, actin, actin-nucleating factors, and prohibitins ‒ IMM scaffolds involved in cristae remodeling. Dephosphorylated BASP1 recruits actin to mitochondria and forms complexes with IMM proteins and α-syn, whereas phosphorylation triggers actin- and Drp1-mediated mitochondrial fission and neurite extension. Constitutive dephosphorylation of BASP1, as occurs under α-syn proteotoxic stress, blocks actin polymerization, disrupts mitochondrial fission and neurite growth, and drives α-syn accumulation in neurites. Together, our findings establish BASP1 as a central regulator of actin dynamics that couples inner and outer mitochondrial membrane to drive fission and promote neurite outgrowth. Moreover, our findings highlight BASP1 as a potential contributor to the pathogenesis of α-synucleinopathies, underscoring its therapeutic relevance.

## RESULTS

### Phosphorylation of human BASP1 at S40 and S170 is regulated by calcineurin and promotes neurite branching in PC12 cells and primary embryonic rat cortical neurons

To identify the substrates dephosphorylated by CaN and associated with α-syn toxic *vs* protective responses we took an unbiased phospho-proteomic approach using isobaric Tandem Mass Tagging (TMT) in combination with mass spectrometry (MS) (*23*). We employed a rat model of PD in which α-syn was unilaterally injected into the substantia nigra pars compacta (*23*). Animals received weekly subcutaneous doses of FK506 or vehicle and were stratified based on terminal brain FK506 concentrations. Strikingly, sub-saturating levels of FK506 (<5 ng/ml) were sufficient to prevent neurodegeneration, as evidenced by improved motor performance and restored striatal dopamine and dopamine transporter levels (*23*). To identify the substrates dephosphorylated by CaN we collected striatal tissue from these animals for TMT-MS analysis. After correction for protein abundance, we detected 51 of 3,526 phosphopeptides that changed phosphorylation significantly when comparing α-syn to control animals. Among these, 50% of the phosphopeptides were significantly hypo-phosphorylated in α-syn animals compared to the controls (Fig. S1A). Given that CaN is a phosphatase highly activated under elevated α-syn levels, we focused our analysis on the hypo-phosphorylated peptides to identify putative CaN-dependent substrates. Five of these phosphopeptides restored their phosphorylation to control levels in α-syn animals treated with sub-saturating doses of FK506. These peptides belonged to only two proteins: GAP-43 (also known as F1, neuromodulin, B-50, G50, and pp46) (*23, 37*) and BASP1 (also known as NAP-22 and CAP-23) (Fig. S1A,B). We identified 3 phosphorylation sites within rat BASP1 that were sensitive to CaN-inhibition: T31, S40, and S165 (corresponding to residues T31, S40, and S170 in human BASP1, respectively) (Fig. S1B).

To determine the functional relevance of the phosphorylation sites identified by our unbiased phosphoproteomic TMT-MS analysis, we used a genetic approach. This well-established approach enables the site-specific dissection of phosphorylation function: substitution with a non-polar residue such as alanine mimics dephosphorylation (phosphoablative mutant), while replacement with a negatively charged residue like aspartic acid mimics phosphorylation (phosphomimetic mutant). We first tested the effect of CaN-dependent BASP1 phosphomutants on neurite growth in PC12 cells, a neuronal-like cell line capable of forming neurites upon expression of proteins such as BASP1 (*38–40*). PC12 cells were co-transfected with green fluorescent protein (GFP) to visualize neurites, and either human wild-type (WT) BASP1 or phosphoablative and phosphomimetic mutants of the phosphosites identified by TMT-MS (T31, S40, and S170). As a negative control, we co-transfected the cells with GFP and an empty vector (hereafter referred to as EV). Cells were imaged 24 hours post-transfection and analyzed for neurite branching. As expected, the BASP1 WT increased neurite branching relative to EV; however, only the phosphomimetic mutants promoted branching beyond that of BASP1 WT (Fig. 1A and Fig. S1C). Importantly, statistically significant differences between the phosphoablative and phosphomimetic mutants were observed only at the S40 and S170 phosphorylation sites. Since the T31 site showed no difference between its phosphoablative and phosphomimetic mutants, we focused on the potential synergistic effects of the S40 and S170 phosphorylation sites by generating double mutants. While the phosphomimetic double mutant did not enhance neurite branching beyond the single mutants, the phosphoablative double mutant significantly impaired BASP1’s ability to promote neurite branching (Fig. 1A and Fig. S1C). We next tested the effect of BASP1 phosphomutants on their ability to modulate neurite length. As expected, BASP1 WT increased neurite length compared to EV (Fig. 1A and Fig. S1C). However, only S40A and the phosphomimetic double mutant showed a difference from BASP1 WT: S40A exhibited slightly shorter neurites, while the phosphomimetic double mutant showed modest neurite elongation. Together, these findings demonstrate that dual phosphorylation of human BASP1 at the CaN-sensitive sites S40 and S170, identified through an unbiased phosphoproteomic screen in an *in vivo* model of α-syn pathology, regulates neurite branching in PC12 cells.

**Fig 1.**
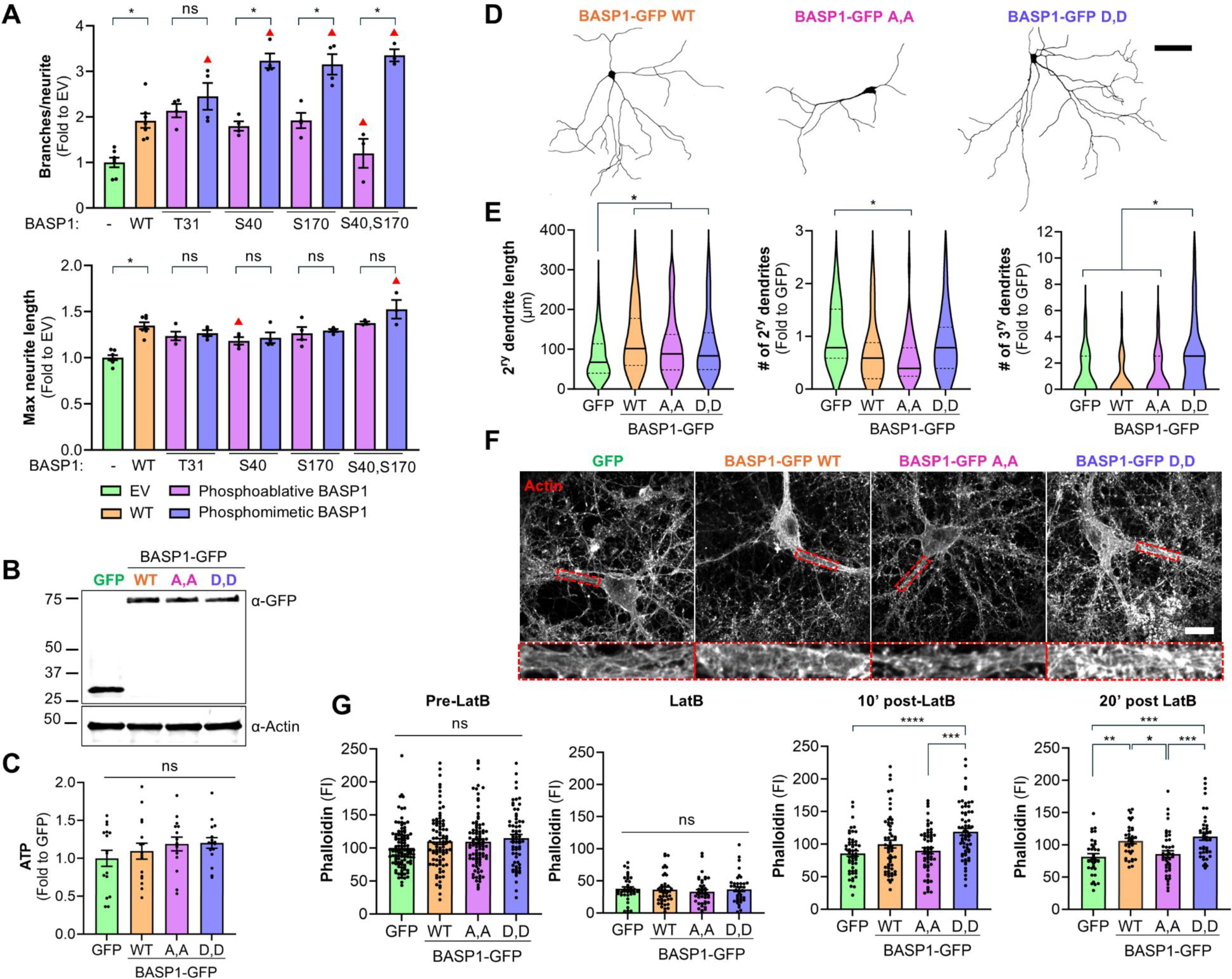
Calcineurin-dependent phosphosites S40 and S170 of human BASP1 control actin polymerization and neurite branching. **(A)** PC12 cells co-transfected with GFP and empty vector (EV) or BASP1, either wild-type (WT) or phosphomutants at the phosphosites identified by MS (T31, S40, and S170) were analyzed for the number of branches and longest neurite length. N>300 cells per condition. One-Way Anova followed by multiple comparisons with two-stage linear step-up procedure of Benjamini, Krieger, and Yekutieli. *q<0.05 comparison between phosphomimetic and phosphoablative mutants, ^▴^q<0.05 comparison between phosphomutants and WT. **(B)** Representative western blot from primary embryonic rat cortical neurons transduced with either GFP alone or BASP1-GFP constructs: WT, phosphoablative (S40A/S170A; A,A), and phosphomimetic (S40D/S170D; D,D) double mutants. Blots were probed with anti-GFP to detect expression of the BASP1-GFP constructs and with anti-actin as a loading control. **(C)** Quantification of ATP levels from conditions in (B). N>30. One-way Anova with post hoc Tukey’s test. **(D)** Representative confocal images of MAP2-immunostained primary embryonic rat cortical neurons cultured under the conditions described in (B). Scale bar = 100 µm. **(E)** Quantification of dendrite complexity in the cultures shown in (D), including measurements of the maximal secondary neurite length, secondary and tertiary dendritic arborization. N≥30. One-Way Anova followed by multiple comparisons with two-stage linear step-up procedure of Benjamini, Krieger, and Yekutieli, *q<0.05. **(F)** Representative confocal images of F-actin stained with BODIPY™ 558/568 Phalloidin in primary embryonic rat cortical neurons cultured under the conditions described in (B). Neurons were treated with 2 µM latrunculin B (LatB) for 10 min, followed by washout and a 10 min recovery period to allow actin re-polymerization. Scale bar = 20 µm. **(J)** Quantification of the levels of F-actin stained with BODIPY™ 558/568 Phalloidin in primary embryonic rat cortical neurons cultured under the conditions described in (B). Mean fluorescent intensities were measured before latrunculin B treatment (Pre-LatB), 10 min after LatB exposure (LatB), and at 10 min and 20 min following LatB washout (10′ and 20′ Post-LatB). N>30. One-way Anova with Tukey’s or Games-Howell’s post-hoc tests (depending on the condition of homoscedasticity and heteroscedasticity), \**P*<0.05, \*\**P*<0.01, \*\*\**P*<0.001, \*\*\*\**P*<0.0001.

We next asked whether phosphorylation of human BASP1 at S40 and S170 also plays a role in neurite branching in neurons. Primary embryonic rat cortical neurons were transduced with lentiviruses expressing either GFP alone as a control or the following GFP-tagged BASP1 constructs: WT, phosphomimetic S40D/S170D double mutant (hereafter referred to as phosphomimetic (D,D) double mutant), and phosphoablative S40A/S170A double mutant (hereafter referred to as phosphoablative (A,A) double mutant). Importantly, none of these mutations affected BASP1-GFP protein expression (Fig. 1B and Fig. S2A) or cell viability, as assessed by ATP levels (Fig. 1C). To visualize the dendritic arbor, we utilized the neuronal specific microtubule-associated protein 2 (MAP2) marker and measured neurite branching by Sholl analysis (Fig. 1D,E). Consistent with our observations in PC12 cells, expression of BASP1-GFP, regardless of phosphomutant status, increased secondary neurite length compared to GFP alone. However, the phosphoablative (A,A) double mutant decreased the number of secondary dendrites compared to control, while the phosphomimetic (D,D) double mutant increased the number of tertiary dendrites compared to both GFP alone and BASP1 WT (Fig. 1D,E).

BASP1 was originally identified as a protein enriched in the cortical cytoskeleton (*30*), and its loss of function has been shown to disrupt the actin cytoskeleton and neurite outgrowth (*33, 39, 40*). To determine whether the effects of BASP1’s CaN-dependent phosphorylation sites on neurite arborization were linked to actin dynamics, we used Latrunculin B (LatB), a pharmacological inhibitor of actin polymerization. LatB binds monomeric (G-) actin with high affinity, preventing its polymerization into filamentous (F-) actin and leading to filament depolymerization. F-actin recovery was assessed at 10, 20, and 30 minutes following LatB washout by staining with BODIPY 558/568 Phalloidin (Fig. 1F). Primary embryonic rat cortical neurons were transduced with GFP alone or GFP-tagged BASP1 WT or double phosphomutants. Prior to LatB treatment, F-actin levels did not differ significantly among the conditions (Fig. 1G and Fig. S3). A 10-minute LatB treatment reduced F-actin levels to ∼50% in all groups (Fig. 1G and Fig. S3). Notably, F-actin recovered the fastest with the phosphomimetic D,D mutant, followed by the WT and then the phosphoablative A,A mutant (Fig. 1G and Fig. S3). Together, these data indicate that the CaN-sensitive sites S40 and S170 on BASP1 regulate actin polymerization and promote dendritic branching in both PC12 cells and primary cortical neurons.

### Phosphorylation of human BASP1 at S40 and S170 mediates interactions enriched in actin- and inner mitochondrial membrane-associated proteins

To identify protein interactors responsible for the BASP1-induced neurite branching, we took an immunoprecipitation (IP) MS-based approach. Primary embryonic rat cortical neurons were transduced with lentiviruses expressing either GFP alone as a control, or GFP-tagged BASP1 constructs: WT, phosphoablative (A,A) and phosphomimetic (D,D) double mutants. IP was performed using GFP-trap agarose beads and analyzed by MS combined with TMT labeling, a quantitative and isobaric labeling which minimizes variability between the samples (Fig. 2A). GFP IPs measured by MS were robust and, importantly, yielded comparable amounts of BASP1 across WT and phosphomutant samples (Table S1). Positive hits were defined as peptides detected in any BASP1-GFP IP (WT and/or phosphomutants) but absent in the GFP-alone IP, based on statistically significant enrichment after normalization to GFP input levels. Using this criterion, we uncovered 185 proteins that specifically bound BASP1 (WT and phosphomutants) (Fig. 2B). By Reactome Pathway analysis, these proteins were enriched for processes involved in endocytosis and neurotransmitter release, functions which have been previously assigned to BASP1 (*40, 41*) (Fig. 2C). To assess how BASP1’s interactors differed between WT and phosphomutants, we normalized spectral counts to BASP1 WT levels (Fig. 2D). Specific hits were defined as those uniquely detected in each condition, showing at least a 1/3-fold change and reaching statistical significance (Table S1). Using these criteria, we identified 23 proteins that specifically bound to the phosphoablative (A,A) double mutant (Fig. 2B,D). By Reactome pathways, these proteins were enriched in two main processes: actin cytoskeleton organization and mitochondrial Ca^2+^-related transport (Fig. 2E). Amongst the actin cytoskeletal components, we identified actin itself (gene name Actb), the actin related proteins 3 and 4 (gene names Actr3 and Arp4), and the actin-folding protein T-complex protein 1 subunit delta (gene name cct4). The mitochondria-enriched proteins included the two prohibitins (gene names Phb1 and Phb2), mitochondrial kinase Phosphoglycerate kinase 1 (gene name PGK1), transforming protein RhoA (gene name Rhoa) and Stomatin-like protein 2 (gene name Stoml2). Interestingly, α-syn was also one of the interactors (gene name SNCA). The phosphomimetic (D,D) double mutant, on the other hand, exhibited only two specific interactors: glycylpeptide N-tetradecanoyltransferase 1 (gene name Nmt1) and cathepsin D (gene name Ctsd). The WT presented a unique interactor, the cell division control protein 42 homolog (gene name Cdc42), a GTPase and key regulator of actin dynamics (*42, 43*). Together, these findings demonstrate that phosphorylation at the CaN-dependent S40 and S170 sites on BASP1 mediates interactions enriched in actin- and mitochondrial-associated proteins.

**Fig 2.**
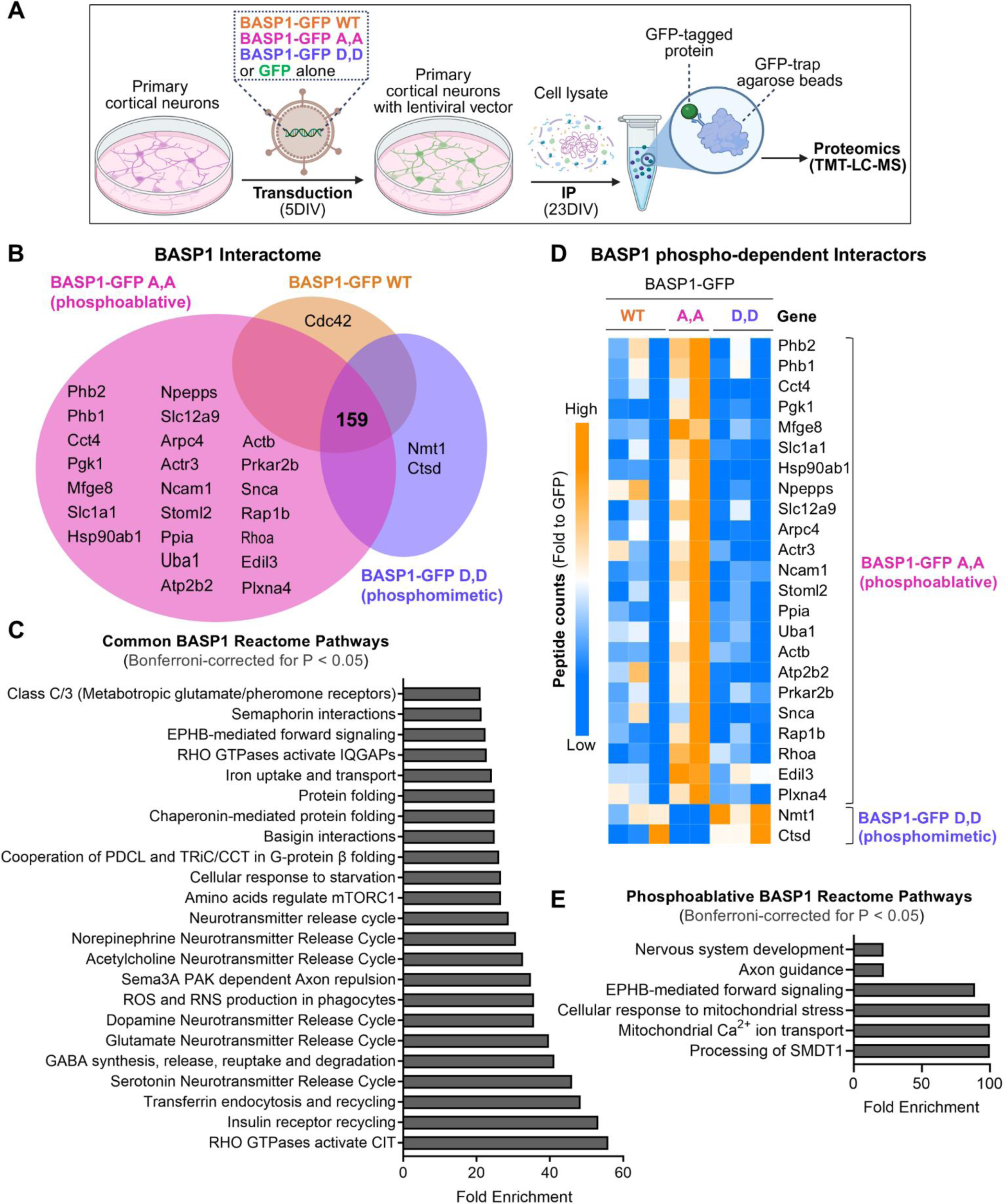
Calcineurin-dependent phosphorylation sites S40 and S170 of BASP1 mediate interactions with actin, actin-nucleating factors, α-synuclein, and prohibitins. **(A)** Schematic representation of the immunoprecipitation (IP) procedure used to isolate BASP1-GFP WT, phosphoablative (S40A/S170A; A,A), and phosphomimetic (S40D/S170D; D,D) double mutant interactors from transduced primary embryonic rat cortical neurons. GFP alone was used as a control. Biorender software was used to create the figure under an academic license. **(B)** Venn diagram showing all mass spectrometry hits from (A) that met the selection criteria detailed in the main text. **(B)** Venn diagram showing all mass spectrometry hits from (A) that met the selection criteria detailed in the main text. **(C)** Reactome pathway analysis for molecular processes for the common BASP1 interactors from (B). **(D)** Heat map representation of CaN-dependent phosphoproteomic hits associated with BASP1 after normalization to BASP1 WT. Each row corresponds to a protein for which a phosphorylated peptide was identified. One-way ANOVA followed by a multiple comparison test with a two-stage linear step-up procedure of Benjamini, Krieger, and Yekutieli. (**E)** Reactome pathway analysis for molecular processes from the CaN-dependent phosphoproteomic hits for BASP1.

### Phosphorylation of human BASP1 at S40 and S170 regulates mitochondrial fission

One of the largest groups of CaN-dependent BASP1 interactors were inner mitochondrial membrane proteins, and amongst those ‒ the two prohibitins (gene names Phb1 and Phb2). To validate prohibitin as a binding partner, we performed IP from HeLa cells transiently expressing either GFP alone or GFP-tagged BASP1 WT or double phosphomutants, followed by western blotting for prohibitin 1. Consistent with our MS data, western blotting showed that BASP1 binds specifically to prohibitin 1, with the phosphoablative (A,A) double mutant exhibiting the strongest interaction (Fig. 3A,B, Table S1). Although prohibitins have been shown to function in the nucleus (*44*) and plasma membrane (*45*), they are best known for their roles in the inner mitochondrial membrane (*46*). In mitochondria, they form large ring-like complexes as heterodimers and play a crucial role in maintaining mitochondrial morphology indirectly modulating dynamics of fission and fusion (*47–50*).

**Fig. 3.**
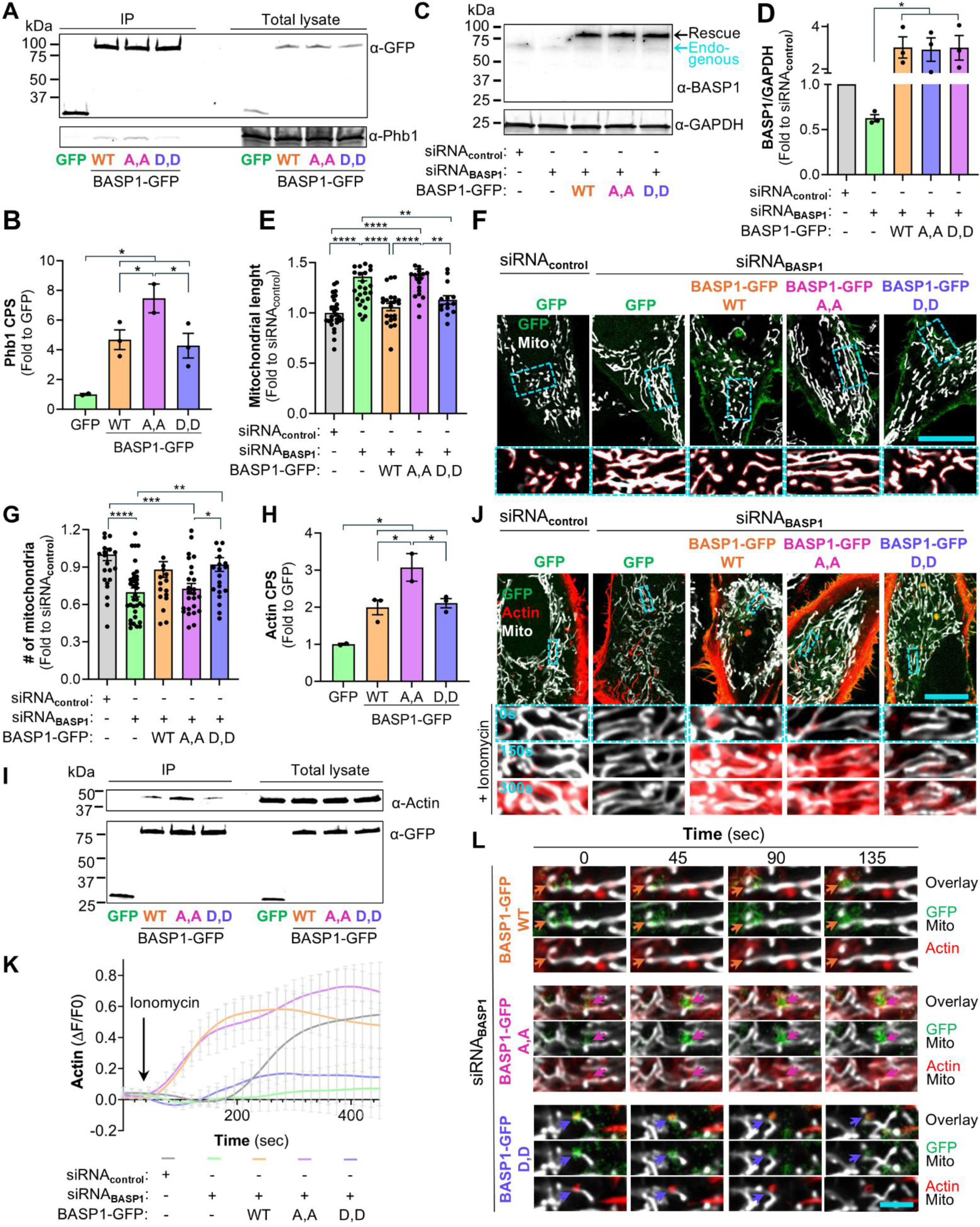
Phosphorylation of human BASP1 at S40 and S170 regulates actin-driven mitochondrial fission in HeLa cells. **(A)** Representative Western blot of GFP immunoprecipitates (IP) from HeLa cells transfected with GFP alone or BASP1-GFP constructs: wild-type (WT), phosphoablative (S40A/S170A; A,A), and phosphomimetic (S40D/S170D; D,D) double mutants. Blots were probed with anti-GFP and anti-prohibitin 1 (Phb1). **(B)** MS quantification of prohibitin 1 (Phb1) co-immunoprecipitated using anti-GFP beads from primary embryonic rat cortical neurons transduced with GFP alone or BASP1-GFP constructs: WT, phosphoablative (S40A/S170A; A,A), and phosphomimetic (S40D/S170D; D,D) double mutants. One-Way Anova followed by multiple comparisons with two-stage linear step-up procedure of Benjamini, Krieger, and Yekutieli, *q<0.05. **(C)** Representative Western blot of BASP1 from HeLa cells co-transfected with GFP alone or BASP1-GFP constructs: BASP1 WT, phosphoablative (S40A/S170A; A,A), and phosphomimetic (S40D/S170D; D,D) double mutants, along with either a 3’UTR-targeting siRNA against human BASP1 or non-targeting control siRNA. **(D)** Quantification of BASP1 signal normalized to GAPDH (loading control) from (C), with values further normalized to the non-targeting siRNA control within each experiment. N=3. One-way Anova with post hoc Dunnett’s test, \**P*<0.05. **(E-G)** Mitochondrial length (E) and number (G) analyzed in HeLa cells from (C) using MitoTracker Deep Red-assisted confocal microscopy (F). Scale bar = 10 µm. N>20. One-way Anova with post hoc Tukey’s test, \**P*<0.05, \*\**P*<0.01, \*\*\**P*<0.001, \*\*\*\**P*<0.0001. **(H)** MS quantification of actin co-immunoprecipitated using anti-GFP beads from primary embryonic rat cortical neurons cultured under the conditions described in (B). One-Way Anova followed by multiple comparisons with two-stage linear step-up procedure of Benjamini, Krieger, and Yekutieli, *q<0.05. **(I)** Representative Western blot of GFP co-immunoprecipitates from HeLa cells cultured under the conditions described in (A). Blots were probed with anti-GFP and anti-actin. **(J)** Representative confocal pictures of mitochodnira and actin from HeLa cells cultured under the conditions described in (C) and co-transfected with LifeAct-mScarlet to visualize actin. Time-lapse imaging was performed at 150-second intervals to monitor actin localization at mitochondria following 4 µM ionomycin stimulation. Scale bar = 10 µm. **(K)** Quantification of actin fluorescence intensity from (J) over time following 4 µM ionomycin treatment. **(L)** Representative confocal pictures of HeLa cells cultured in conditions described in (C) under 4 µM Ionomycin treatment. Actin was visualized by transient expression of LifeAct-mScarlet and mitochondria were labeled with MitoTracker Deep Red. Time-lapse imaging was performed at 15-second intervals. Arrows indicate actin recruitment to mitochondria with BASP1-GFP WT, phosphoablative (S40A/S170A; A,A), and phosphomimetic (S40D/S170D; D,D) double mutants followed by fission events only in BASP1 WT and D,D. Scale bar = 2 µm.

Given that prohibitins are highly expressed in mitochondria and that the BASP1 phosphoablative mutant preferentially interacts with these and other inner mitochondrial membrane proteins, we investigated whether BASP1 contributes to the regulation of mitochondrial morphology. To evaluate the role of endogenous BASP1 and the phosphomutants, we first employed HeLa cells. We co-transfected HeLa cells with an siRNA against the 3’UTR of human BASP1 to knockdown BASP1 or siRNA encoding a scrambled sequence as a control, with either GFP alone, or GFP-tagged BASP1 (WT, phosphoablative (A,A) and phosphomimetic (D,D) double mutants) to track and complement for the lack of endogenous BASP1 (Fig. 3C,D). Mitochondria were visualized with MitoTracker Deep Red followed by morphometric analysis using ImageJ/Fiji (Fig. 3E-G). Knockdown of BASP1 significantly increased mitochondrial length compared to control (Fig. 3E,F). Importantly, this effect was specific to BASP1, as complementation with BASP1 WT rescued the mitochondrial length defects caused by the absence of BASP1 (Fig. 3E,F). Complementation with the phosphoablative (A,A) double mutant however, did not rescue the mitochondrial length defects caused by reduction of endogenous BASP1, whereas the phosphomimetic (D,D) double mutant did (Fig. 3E,F). In agreement with the defects in mitochondrial length caused by BASP1 knockdown, it also significantly decreased the number of mitochondria compared to control (Fig. 3F,G). Importantly, the reduction in mitochondrial number was rescued by expression of the phosphomimetic (D,D) double mutant, but not by the phosphoablative (A,A) double mutant (Fig. 3F,G). Together, these data indicate that BASP1 plays a key regulatory role in mitochondrial morphology.

### Phosphorylation of human BASP1 at S40 and S170 regulates actin-driven mitochondrial dynamics

Mitochondrial morphology is dynamically shaped by the balance between fusion and fission processes, which are tightly regulated by several proteins, cellular signals, and bioenergetic status (*51*). Actin polymerization plays a crucial regulatory and mechanical role in mitochondrial fission (*2, 6, 8–17*). Moreover, mitochondrial fission depends on *de novo* actin polymerization at mitochondria (*10, 52*). We showed that BASP1 regulates actin polymerization (Fig. 1F,G). Further, IP-MS analysis revealed that BASP1 binds actin, with the phosphoablative (A,A) double mutant showing the strongest binding compared to both the WT and the phosphomimetic (D,D) double mutant (Fig. 3H, Table S1). We therefore investigated whether BASP1 regulates mitochondrial morphology through actin polymerization, thereby promoting mitochondrial fission. To validate actin as a binding partner, we performed IP followed by western blotting for actin from HeLa cells transiently expressing GFP alone or GFP fusions of BASP1WT or double phosphomutants. Consistent with the MS data, the phosphoablative (A,A) double mutant exhibited the highest actin binding (Fig. 3I).

We next asked whether phosphorylation of human BASP1 at the S40 and S170 sites directly contributes to mitochondrial actin dynamics. To test this, we performed time-lapse imaging of mitochondria and actin in HeLa cells co-transfected with: 1) an siRNA targeting the 3′UTR of human BASP1 to deplete endogenous BASP1, or with scrambled siRNA as a control; 2) either GFP alone or GFP-tagged BASP1 constructs (WT, phosphoablative (A,A), and phosphomimetic (D,D) double mutants) to assess functional rescue; and 3) LifeAct-mScarlet, a fluorescent probe that binds both monomeric and filamentous actin. Mitochondria were visualized with MitoTracker Deep Red, and cytosolic Ca^2+^ influx was induced using ionomycin. In control cells, ionomycin treatment induced a progressive increase in actin accumulation around mitochondria, a response that was abolished by BASP1 knockdown (Fig. 3J,K and Fig. S2B). This phenotype was specific to BASP1, as re-expression of BASP1 WT fully restored ionomycin-induced actin recruitment to mitochondria. The phosphoablative (A,A) double mutant also rescued actin accumulation to BASP1 WT levels, whereas the phosphomimetic (D,D) double mutant showed minimal actin recruitment (Fig. 3J,K and Fig. S2B). Collectively, BASP1 knockdown results in elongated mitochondria (Fig. 3E) and loss of peri-mitochondrial actin enrichment (Fig. 3K). Similarly, the phosphoablative (A,A) double mutant results in elongated mitochondria, but unlike the knockdown condition, it exhibited elevated actin accumulation at mitochondria. In contrast, the phosphomimetic (D,D) mutant results in shorter mitochondria and reduced actin occupancy at mitochondria. These findings suggest that BASP1 phosphorylation at CaN-dependent sites regulates actin recruitment to mitochondria: dephosphorylated BASP1 promotes actin binding, while phosphorylation may be required to release actin for polymerization and enable fission. To test this, we performed live-cell imaging under ionomycin-induced Ca^2+^ influx conditions with 15-sec intervals (Fig. 3L and Movies S1-S3). Upon Ca^2+^ stimulation, BASP1 WT and actin localized to mitochondria and promoted fission (Fig. 3L and Movie S1). The phosphoablative (A,A) double mutant BASP1 recruited actin to mitochondria but failed to support fission (Fig. 3L and Movie S2). In contrast, the phosphomimetic (D,D) double mutant BASP1 recruited less actin yet promoted fission (Fig. 3L and Movie S3). Collectively, these data demonstrate that Ca^2+^-dependent BASP1 phosphorylation dynamically regulates actin engagement at mitochondria to drive fission, and that disruption of this phospho-cycle impairs actin turnover and fission kinetics.

### BASP1 modulates mitochondrial fission by controlling Drp1 recruitment

Actin polymerization spatially defines fission sites and facilitates the recruitment and function of Drp1, a GTPase essential for mitochondrial constriction and division (*12, 53*). If Ca^2+^-dependent regulation of BASP1 is required to initiate actin recruitment at mitochondria to drive fission, then BASP1 should act upstream of Drp1. To test this hypothesis, HeLa cells were co-transfected with Drp1-mCherry along with the following constructs: 1) an siRNA targeting the 3′UTR of human BASP1 to deplete endogenous BASP1, or with scrambled siRNA as a control, and 2) either GFP alone or GFP-tagged BASP1 constructs (WT, phosphoablative (A,A), and phosphomimetic (D,D) double mutants) to assess functional rescue. Mitochondria were visualized with MitoTracker Deep Red. As expected, BASP1 knockdown abolished Drp1-mCherry clustering at mitochondria causing a decrease in both the number and size of Drp1 clusters compared to the control (Fig. 4A-C). We observed Drp1-mCherry clusters enclosed within well-defined, donut-shaped BASP1-GFP structures (Fig. 4A). Drp1 clustering depended on BASP1 regardless of its phosphorylation state, as complementation with BASP1 WT, phosphoablative (A,A), and phosphomimetic (D,D) double mutants restored Drp1 recruitment and clustering at mitochondria (Fig. 4A–C). However, only WT (Fig. 4D and Movie S4) and the phosphomimetic D,D mutant (Fig. 4D and Movie S5) supported fission under ionomycin-induced Ca^2+^ stimulation, whereas the A,A mutant did not (Fig. 4D and Movie 6). Together, these findings demonstrate that BASP1 is necessary for Drp1 clustering at mitochondria and that CaN-dependent regulation of BASP1 is required for Drp1 activity and mitochondrial fission.

**Fig. 4.**
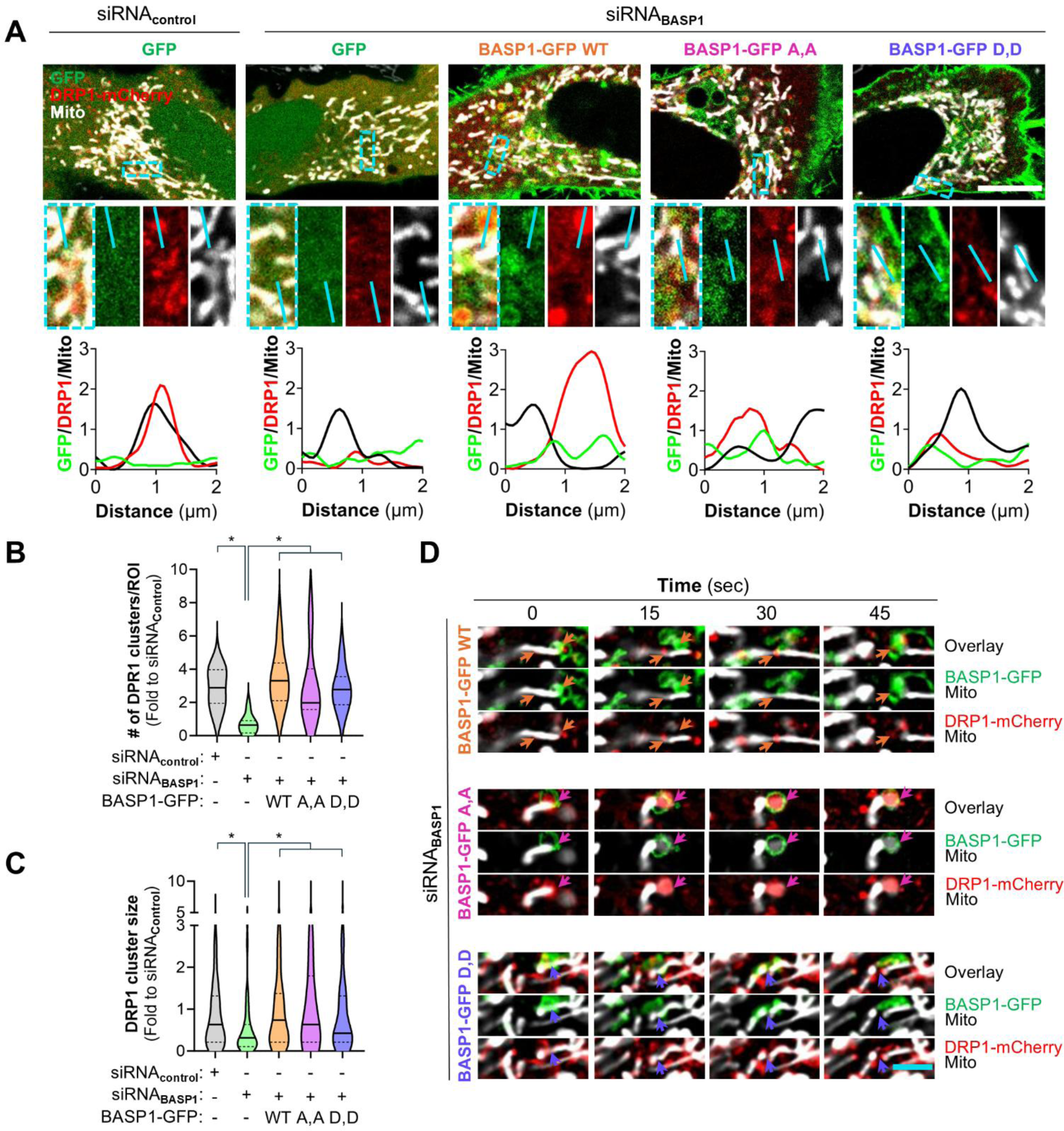
BASP1 regulates Drp1 recruitment, and calcineurin-dependent phosphorylation at S40 and S170 is required for Drp1-mediated mitochondrial fission. **(A)** Representative confocal microscopy images of HeLa cells co-transfected with siRNA (control or targeting 3’UTR of human BASP1) and either GFP alone or BASP1-GFP constructs: wild-type (WT), phosphoablative (S40A/S170A; A,A), and phosphomimetic (S40D/S170D; D,D) double mutants. Drp1 was visualized by transient expression of Drp1-mCherry and mitochondria were labeled with MitoTracker Deep Red. Blue lines correspond to line-scans across Drp1 clusters. Scale bar = 10 µm. Number per ROI **(B)** and size **(C)** of Drp1 clusters were quantified in HeLa cells from (A). N>150. One-Way Anova followed by multiple comparisons with two-stage linear step-up procedure of Benjamini, Krieger, and Yekutieli, *q<0.05. **(D)** Representative confocal images of Drp1-mCherry recruitment to MitoTracker Deep Red-stained mitochondria in HeLa cells from (A) under 4 µM Ionomycin treatment (15 sec intervals). Scale bar = 2 µm.

### Phosphorylation of human BASP1 at S40 and S170 regulates mitochondrial dynamics, α-synuclein aggregation, and neurite outgrowth

To assess whether CaN-dependent phosphorylation of BASP1 regulates actin-mediated mitochondrial fission in neurons, we employed an *in vitro* scratch assay, also known as a neurite regeneration assay, for three key reasons. First, it provides a well-defined and reproducible model of neuronal injury, where regeneration is driven by Ca^2+^-dependent actin polymerization at growth cone (*54, 55*), facilitating mitochondrial remodeling and microtubule outgrowth (*56*). Second, expression of BASP1 phosphomutants regulates neurite arborization (Fig. 1E) and modulates actin polymerization (Fig. 1G) in primary embryonic rat cortical neurons. Third, *in vivo* data from the rat model of α-syn pathology revealed that BASP1 phosphorylation state correlates with neurite integrity: dephosphorylated BASP1 associates with neurite degeneration, whereas phosphorylated BASP1 correlates with neurite protection and structural preservation (Fig. S1A,B and (*23*)). Primary embryonic rat cortical neurons transduced with GFP alone or GFP-tagged BASP1 WT or double phosphomutants were mechanically injured on day 22 *in vitro* (DIV22) and allowed to regenerate their neurites over a 24-hour period. To visualize the newly formed neurite processes we immunostained the cultures for the neuron-specific microtubule protein βIII-tubulin. Consistent with our findings on actin polymerization (Fig. 1F,G), the BASP1 phosphomimetic (D,D) double mutant exhibited the fastest neurite regrowth (Fig. 5A,B and Fig. S4).

**Fig. 5.**
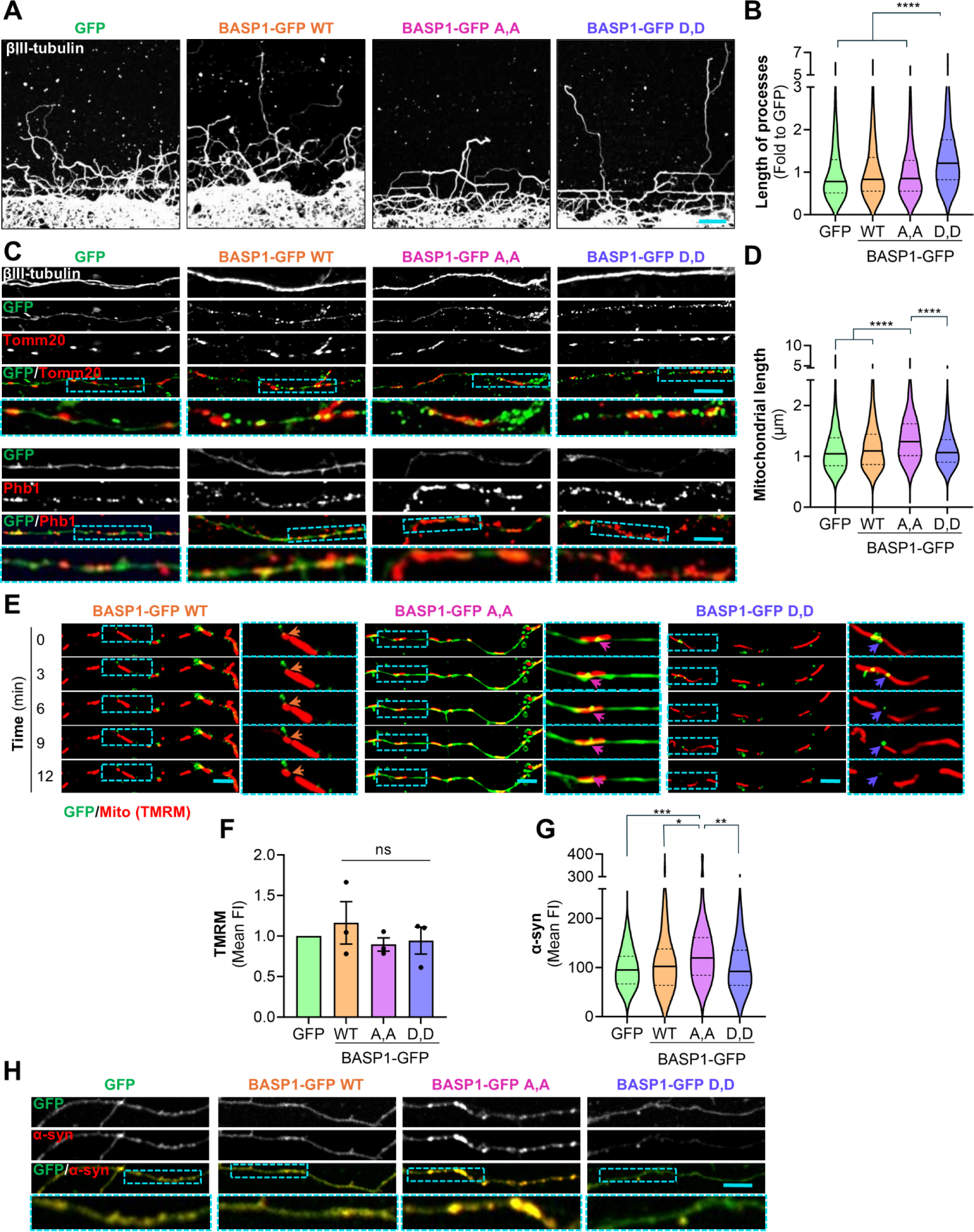
Calcineurin-sensitive phosphorylation sites on BASP1 regulate mitochondrial dynamics to promote neurite outgrowth in primary embryonic rat cortical neurons. **(A)** Representative confocal images of β-III-tubulin staining in DIV23 primary embryonic rat cortical neurons, transduced with either GFP alone or BASP1-GFP constructs: wild-type (WT), phosphoablative (S40A/S170A; A,A), and phosphomimetic (S40D/S170D; D,D) double mutants. Neurons were subjected to mechanical injury in a scratch assay, and the staining was performed 24 hours post-injury. Scale bar = 50 µm. **(B)** Quantification of neurite outgrowth for the cultures in (A) using ImageJ/Fiji. N>950. One-Way Anova with Games-Howell’s post-hoc test, \*\*\*\**P*<0.0001 **(C)** Representative immunofluorescence images of primary embryonic rat cortical neurons cultured under the conditions described in (A) showing immunostaining for mitochondrial markers TOMM20 and prohibitin 1 (Phb1). Scale bar = 5 µm. **(D)** Quantification of mitochondrial length in neurites for the cultures in (C) N>550. One-way Anova with Games-Howell’s post-hoc test, \*\*\*\**P*<0.0001. **(E)** Representative live-cell images of primary embryonic rat cortical neurons cultured under the conditions described in (A) and stained with the fluorescent mitochondrial dye tetramethylrhodamine methyl ester (TMRM) acquired at 3-min intervals. Scale bar = 5 µm. **(F)** Quantification of TMRM intensities for the cultures in (E), normalized to GFP alone within each experiment. N=3. One-way Anova with post hoc Tukey’s test. **(G)** Quantification of α-syn in neurites from neurons cultured under the conditions described in (A). N>80. One-way Anova with Games-Howell’s post-hoc test, \**P*<0.05, \*\**P*<0.01, \*\*\**P*<0.001. **(H)** Representative immunofluorescence images of primary embryonic rat cortical neurons cultured under the conditions described in (A) showing staining for α-synuclein (α-syn). Scale bar = 5 µm.

To investigate whether the effects on neurite outgrowth were associated with changes in mitochondrial morphology, we immunostained the mechanically injured neuronal cultures with TOMM20, OMM protein, and Phb-1, IMM protein and BASP1 phospho-specific interactor (Fig. 3B). Consistent with our findings in HeLa cells, the mitochondria were significantly elongated in the presence of the phosphoablative (A,A) double mutant compared to both GFP alone control and BASP1 WT (Fig. 5C,D).

To assess whether these mitochondrial defects resulted from impaired dynamics, we performed live-cell imaging with TMRM, a membrane potential–sensitive fluorescent dye, while monitoring BASP1 localization via its GFP tag. As expected, BASP1 WT supported mitochondrial fission in regenerating neurite processes (Fig. 5E, Movie S7). Although, none of these mutations affected the mitochondrial membrane potential as determined by TMRM intensity (Fig. 5F), we only detected fission events in the presence of the BASP1 WT and phosphomimetic (D,D) double mutant but not in the phosphoablative (A,A) double mutant, in the time frame recorded (Fig. 5E and Movie S7).

Finally, we examined α-syn intracellular localization, given its preferential binding to the phosphoablative (A,A) BASP1 mutant in our IP-MS analysis (Fig. S2C and Table S1). Immunostaining revealed increased accumulation of endogenous α-syn in regenerating neurite processes expressing phosphoablative (A,A) double mutant when compared to GFP alone, BASP1 WT and the phosphomimetic (D,D) double mutant (Fig. 5G,H). Collectively, these findings demonstrate that phosphorylation at S40 and S170 is essential for BASP1-mediated mitochondrial fission in developing neurites. Disruption of this phosphorylation equilibrium impairs fission, promotes α-syn accumulation, and ultimately compromises neurite regrowth.

## DISCUSSION

Although actin is a critical driver of mitochondrial fission (*2, 6, 8–17*), the mechanisms linking actin remodeling to mitochondrial division and coordinating constriction of the inner and outer membranes remain poorly understood. Our findings reveal that BASP1 acts as a dynamic molecular switch connecting Ca^2+^ signaling to mitochondrial fission. Under resting conditions, BASP1 is predominantly phosphorylated. Cytosolic Ca^2+^ elevation activates CaN, which dephosphorylates BASP1 and promotes its interaction with actin, α-syn and inner mitochondrial membrane proteins. This recruitment primes the fission machinery, and subsequent BASP1 rephosphorylation triggers actin- and Drp1-mediated mitochondrial fission, thus coordinating division of the inner and outer membranes. Newly divided mitochondria, together with actin polymerization at growth cones, support neurite outgrowth by promoting microtubule assembly and the inheritance of healthy mitochondria to nascent neuronal processes (Fig. S5A). In contrast, constitutive BASP1 dephosphorylation, as observed under α-syn–induced proteotoxic stress, increases its binding to actin, α-syn, prohibitins, and other inner mitochondrial membrane proteins. This aberrant interaction impairs mitochondrial fission, resulting in elongated mitochondria, α-syn accumulation, and the formation of less complex neurites (Fig. S5B).

The actin cytoskeleton is known to play a key role in mitochondrial fission at the OMM, in coordination with ER–mitochondria contact sites and OMM-anchored adaptor proteins that recruit Drp1 (*6, 13, 57, 58*). While it has been proposed that the IMM undergoes remodeling following Drp1-driven OMM constriction, this step is less well understood and is thought to depend on mechanical force transmitted from the OMM. Our data suggests that BASP1-mediated actin polymerization physically bridges the IMM and OMM in a process regulated by CaN activity providing the missing link.

Proper mitochondrial dynamics is essential for cellular function, and their disruption is linked to numerous human diseases (*59*). Defects in mitochondrial function and actin cytoskeleton organization have long been implicated in α-syn pathobiology (*60, 61*). BASP1 has been bioinformatically identified as a critical hub in Parkinson’s disease (*36*), yet its contribution to pathology remains unclear. Phosphoproteomic analysis in an α-synucleinopathy model revealed BASP1 as a Ca^2+^/CaN-regulated mediator of α-syn pathology. Pathological Ca^2+^ elevation drives sustained BASP1 dephosphorylation, trapping actin at the inner mitochondrial membrane, inhibiting fission, and promoting α-syn accumulation in neurites, establishing a feedforward loop that accelerates neurodegeneration (Fig. S5B). BASP1 phosphorylation thus emerges as a central node linking actin remodeling to mitochondrial dynamics, positioning BASP1 as a potential therapeutic target in α-synucleinopathies.

## MATERIALS AND METHODS

### Plasmids and viruses

The BASP1 mutagenesis was performed using Q5 Site-Directed Mutagenesis Kit (E0554, New England Biolabs) according to the manufacturer protocol. Human BASP1 pcDNA3.1-Myc-His construct was used as a template DNA. T31, S40 and S170 were substituted with either A or DD. For the experiments with primary embryonic rat cortical neurons, the phosphoablative S40A,S170A and phosphomimetic S40D,S170D double mutants were obtained by mutagenesis of BASP1 WT subcloned into the lentiviral expression vector pER4-eGFP. Viruses were produced in the NU Gene editing transduction and Nanotechnology Core. Equal MOIs were used to infect rat cortical cultures on DIV5.

mCh-Drp1 (Addgene, 49152) and pLifeAct-mScarlet3_N1 (Addgene, 189767) were employed for live-cell imaging.

Non targeting ON-TARGETplus Control siRNA (Dharmacon, D-001810-01-20) and targeting 3’UTR of human BASP1 siGENOME siRNA (Dharmacon, D-019008-19-0020) were purchased from Dharmacon.

### Cell lines

HeLa cells were maintained in Dulbecco’s modified Eagle’s medium (DMEM, Gibco, 11965092) supplemented with 10% fetal bovine serum (FBS, Denville) in humidified 5% CO_2_ incubator at 37°C. Cells were released from culture dishes by treatment with 0.05% trypsin and 2 mM EDTA (Gibco, 15050-065) in phosphate-buffered saline (PBS) at 37°C for 5 min. Cell viability was evaluated by the trypan blue (0.4%, Gibco 15250-061) exclusion test. Lipid-mediated transfection of HeLa cells was performed using Lipofectamine 2000 (Thermofisher, L3000015). In details, 2.5 µg of DNA and 3 µL of 20 µM siRNA were added to 150 µL Opti-MEM media (Gibco, 31985-062) in a sterile Eppendorf tube. 5 µL of Lipofectamine 2000 were added to 150 µL Opti-MEM media in separate tube. Following 5 min incubation, the solutions were combined for another 20 min incubation at 25 °C and added to a 35 mm FluoroDish (World Precision Instrument FD35100) with pre-cultured HeLa cells in DMEM (Gibco, 11965092) supplemented with 3% FBS (Denville). 5 hours post-transfection, the media was replaced with DMEM supplemented with 10% FBS (Denville) for another 24 h.

PC12 cells were maintained up to passage 10 in DMEM (Gibco, 11965092) supplemented with 10% NuSerum and 1% FBS (Denville) in humidified 5% CO_2_ incubator at 37°C. 70-80% confluent culture was passaged by trypsinization (0.05% Trypsin/EDTA, Gibco, 15050-065) followed by centrifugation at 500×g for 5 min and resuspension in full culture medium. For the evaluation of the response to BASP1 overexpression, the cells were seeded on collagen-coated tissue culture polystyrene. 1 ml aqueous solution containing 35 µg/ml human placenta collagen IV (Advanced Biomatrix) and 15 µg/ml chicken collagen II (Sigma) was added to each well of a 24-well plate followed by overnight air-drying at 25 °C and UV-sterilization. Lipofectamine 3000 (Invitrogen, L3000008)-assisted transfection was performed 24 h prior to examination. For a single well of 24-well plate, 500 ng pcDNA 3.1/Hygro construct (BASP1-Myc-His WT, phosphoablative and phosphomimetic mutants or EV) and 83 ng pEGFP-C1 were added to 50 µL Opti-MEM (Gibco, 31985-062) along with 0.8 µL P3000 reagent. 0.8 µL of Lipofectamine 3000 reagent was diluted in 50 µL Opti-MEM in a separate sterile Eppendorf tube. Ater 5 min incubation, the DNA and Lipofectamine 3000 reagent solutions were combined, incubated for 30 min and then added to the PC-12 cells pre-cultured for 24 h. After 5 h, the solution was replaced with the full growth media. Cells were imaged using Leica DMI3000B microscope coupled with an QImaging QIClick CCD Camera and Leica HCX PL FLUOTAR L 20×/0.40 CORR PH1 objective. Image acquisition was done using Q-capture pro7 software and manual tracking of equal exposure and digital gain setting between images. Image processing and analysis was done using ImageJ/Fiji.

### Primary cortical cultures

Primary embryonic rat cortical neurons were isolated from euthanized pregnant Sprague–Dawley rats at embryonic day 18 as described elsewhere (*37*). Protocol was approved by Northwestern University administrative panel on laboratory animal care. Embryos were harvested by Cesarean section and cerebral cortices were isolated and dissociated by 0.25% Trypsin (Invitrogen, 15090-046) digestion for 15 min at 37°C followed by washing and trituration with 1 ml plastic tip. After passing through a 70 µm cell strainer (Falcon, 352350), the cells were seeded on tissue Poly-D-Lysine (Sigma, P-1149)-coated 96-well (4 × 10^4^ cells per well), 24-well plates with inserted 12 mm Neuvitro GG-12-1.5-oz coverslips (5 × 10^4^ cells per well), a FluoroDish (3 × 10^5^) or 10 cm dish (4 × 10^6^ cells per dish) in neurobasal medium (Invitrogen, 21103-049) supplemented with 10% heat-inactivated FBS (Denville), 2 mM GlutaMAX Supplement (Gibco, 35050061), penicillin (100 IU/mL), and streptomycin (100 μg/mL) (Gibco, 15140-122). Prior to seeding, the cells were counted using an Automated cell counter TC10 (Bio-Rad) and viability (90-95%) was checked with trypan blue (0.4%, Gibco 15250-061) exclusion test. After 1 h incubation at 37°C, media was changed to neurobasal medium (Invitrogen, 21103-049) supplemented with B27 (Gibco 17504-044) and 2 mM GlutaMAX Supplement (Gibco 35050061). One-half of the media was changed twice a week and on DIV5, 4 h before transduction. As a surrogate marker of cell viability, cellular ATP content was measured using the ATPlite kit (PerkinElmer, 6016941) according to the manufacturer’s instructions.

### Immunoprecipitation (IP)

DIV23 primary embryonic rat cortical neurons transduced with pER4-GFP and pER4 BASP1-GFP (WT and double phosphomutants S40A,S170A and S40D,S170D) were briefly washed with PBS supplied with 0.2mM CaCl_2_ and 1mM MgCl_2_ and lysed using ice-cold lysis buffer [1% Triton X-100, 10% glycerol, 10 mM TrisHCl pH 7.5, 150 mM NaCl, 0.5 mM EDTA] supplemented with the Halt protease and phosphatase inhibitor cocktail (Thermofisher, 78441). Samples were incubated on ice for 30 minutes and pushed through a 27G needle (10 times) to ensure full lysis and then centrifuged at 21,000×g for 20 minutes. Protein concentrations were analyzed with the Pierce BCA Protein Assay kit (Thermo Scientific, PI23227) and Fisherbran accuSkan GO UV/Vis Microplate Spectrophotometer (Fisher Scientific). Each supernatant was incubated rotating at 4°C for 2 h with 30 ul of 50% GFP-Trap beads slurry (ChromoTek GFP-Trap Agarose beads) prepared in lysis buffer. Next, the beads were washed trice in a centrifuge (2,500×g, 2 min, 4°C) with lysis buffer and two more times – with the wash buffer [10mM Tris-HCl pH7.5, 150mM NaCl, 0.5mM EDTA]. The first elution was performed by 30 min incubation in Elution buffer I [50mM Tris-HCl pH7.5, 2 M Urea, 5µM Sequencing Grade Modified Trypsin, (Promega), 1mM DDT] at 30°C with agitation (4000 rpm) followed by centrifugation at 2,500×g for 2 min. The second eluate was collected in Elution buffer II [50mM Tris-HCl pH7.5, 2 M Urea, 5mM Iodoacetamide]. The eluates were combined and submitted for TMT-MS at the MIT Kock Proteomics facility.

IP from HeLa cells transiently expressing pER4-GFP and pER4 BASP1-GFP (WT and double phosphomutants S40A,S170A and S40D,S170D) were analyzed by western blotting after the agarose bead washing step.

### Western blotting

Transfected HeLa cells were rinsed with PBS supplied with 0.2mM CaCl_2_ and 1mM MgCl_2_ and then lysed with ice-cold IP buffer supplemented with DNAse (Millipore Sigma, 101041590001) and Halt protease and phosphatase inhibitor cocktail (Thermofisher, 78441) for 20 min with subsequent addition of sodium dodecyl sulfate (Sigma) to the final concentration of 1% for another 20 min. Samples were pushed through a 27G needle (10 times) to ensure full lysis and then centrifuged at 21,000×g for 20 minutes. The obtained supernatants were used for Western blotting analysis. Protein concentration was analyzed with the Pierce BCA Protein Assay kit (Thermo Scientific, PI23227) and Fisherbrand accuSkan GO UV/Vis Microplate Spectrophotometer (Fisher Scientific). After the boiling with an appropriate amount of the 6× Laemmli Sample Buffer (Bioland scientific LLC, sab03-02) and 5% β-mercaptoethanol (Sigma, 444203), the protein samples (∼30 µg) were separated on precasted 4-20% Criterion TGX Stain-free gels (Bio-Rad) and transferred to nitrocellulose membranes (Amersham Protran 0.2um NC, #10600001). Membranes were then blocked with 3% bovine serum albumin (BSA, Sigma, SLBT8252) in Tris-buffered saline (TBS, 50 mM Tris, pH 7.4, 150 mM NaCl) with 0.1% Tween (TBST) for 1 h at 25°C and immunoblotted overnight in primary antibodies (anti-BASP1, Biomatik, CAU32091; anti-β-Actin, Abcam, ab6276; anti-Prohibitin, Abcam, ab75771; anti-Na^+^/K^+^ ATPase alpha-3, Sigma, 06-172; Anti-GAPDH, Proteintech, 60004; anti-GFP, Abcam, ab183734; anti-GFP, Santa Cruz, sc-9996; anti-H3, Cell Signalling, 14932) in the blocking buffer, at 4°C, shaking. The following day, the membranes were washed three times with TBST for 5 minutes each and incubated with secondary antibodies (IRDye or goat anti-mouse IgG H&L-HRP, Abcam, ab6789) for 1 h shaking at 25°C followed by three washes with TBST and imaging using Li-Cor Odyssey® CLx Imaging System or Biorad ChemiDoc MP Imaging System, accordingly. Images were processed and analyzed using ImageJ/Fiji (*62*).

### Scratch assay

In order to evaluate the neurite regeneration propensity following injury, we scratched the transduced DIV22 primary embryonic rat cortical cultures across the diameter of the coverslip by a disposable sterile tip (Eppendorf, 200 µL). Then, we incubated the cells for 24 h and evaluated the neurite outgrowth by immunostaining for β-III-tubulin, mitochondrial morphology – by immunostaining for TOMM20 and Phb1, α-syn accumulation – by immunostaining for α-syn.

### Immunofluorescence

Transduced neuronal cultures (DIV23) were rinsed with PBS supplied with 0.2mM CaCl_2_ and 1mM MgCl_2_ and then fixed with 4% (vol/vol) paraformaldehyde (PFA, Polysciences, 04018-1) in PBS with 4% Sucrose (Sigma, S7903). Fixed cells were washed trice with PBS and permeabilized using 0.15% Triton X-100 in PBS, at 25°C for 10 min. Permeabilized cells were washed trice with PBS and blocked in 10% normal goat serum (Invitrogen, PCN5000) in PBS with 0.1% Tween (PBST) for 60 min at RT. Primary antibodies (anti-MAP2, Millipore, AB5622; anti-MAP2, Abcam, ab5392; anti-βIII-tubulin, NB100-1612; anti-βIII-tubulin, Invitrogen, MA1-118; anti-TOMM20 [4F3], Abcam, ab56783; anti-Prohibitin, Abcam, ab75771; anti-α-syn [LB509], Abcam, ab27766; anti-GFP, Abcam, ab183734; anti-GFP, Invitrogen, A6455) were prepared in 2% normalized goat serum and 1% BSA (Sigma, A7906) in PBST. The cells were incubated with primary antibodies overnight at 4°C followed by triple wash and incubation (1 h at 25°C) with secondary antibodies prepared in the same buffer. Subsequently, the cells were stained with Hoechst (Invitrogen 33342) for 5 min at 25°C followed by triple rinsing with PBS and mounting with ProLong Gold Antifade Mountant (Invitrogen, P36930). Cells were imaged using Leica MI4000B confocal microscope with ACS APO 63x/1.30 OIL objective and Leica LasX software at 1024×1024 pixel resolution. Images were processed and analyzed using ImageJ/Fiji (*62*).

### Latrunculin B washout

Transduced neuronal cultures (DIV23) were incubated with 2µM Latrunculin B (Sigma-Aldrich, L5288) for 10 min followed by washout with full media for 10-30 min. F-actin was visualized by BODIPY 558/568 Phalloidin staining. The neurons were fixed with 4% PFA (Polysciences, 04018-1) in PBS with 4% Sucrose (Sigma, S7903) for 15 minutes at 25°C, then washed with PBS trice for 5 min each and permeabilized with 0.1% Triton X-100 in PBS for another 15 minutes at RT. Next, the cells were washed twice with PBS and stained for 1 h at 25°C in the dark with 165nM BODIPY 558/568 Phalloidin prepared in PBS with 1% BSA (Sigma, A7906). Finally, the cells were washed with PBS trice for 5 min each, immunostained for MAP2, and mounted on coverslips.

### Sholl analysis

Sholl analysis was performed using the SNT plugin (ImageJ/Fiji) after manual tracing of DIV23 primary embryonic rat cortical neurons visualized by MAP2 immunostaining. The neurons were imaged using BZX optical microscope (Keyence, Osaka, Japan) with 20× Objective Lens, Advanced Observation Module (XY-stitching, z-stacking, multi-point acquisition) and Optical Sectioning Module (Structured Illumination Microscopy).

### Live-cell imaging

Live-cell imaging was performed with Nikon AXR NSPARC point-scanning laser confocal microscope using 100× (NA=1.49), at 2048×2048 pixel resolution at Center for Advanced Microscopy at Nikon Imaging Center at Northwestern University. HeLa cells were used for imaging Ca^2+^ influx-triggered mitochondrial and actin dynamics by means of staining with MitoTracker Deep Red FM (Cell Signalling, 8778S) and transfection with pLifeAct-mScarlet3_N1 (Addgene, 189767), respectively. Drp1 recruitment was visualized using mCh-Drp1 (Addgene, 49152) construct. Images were acquired with 15 sec intervals in Phenol-free DMEM (Gibco, A14430-01), supplemented with 1g/l glucose (Sigma-Aldrich, G8769) and 10% FBS (Denville). Ionomycin (Invitrogen,124222) was added to the cells to the final concentration of 4 µM at 45 sec timepoint. Actin filaments were measured within regions of interest (ROIs) that excluded stress fibers and cell edges. NIS-Elements software was used for image acquisition and processing with Denoise.ai and Deconvolution tools. Image analysis was performed using ImageJ/Fiji and CellProfiler.

40 nM TMRM (Invitrogen, T668) was used for visualization of mitochondrial dynamics in pER4 BASP1-GFP-transduced (WT and double phosphomutants S40A,S170A and S40D,S170D) primary embryonic rat cortical neurons in Phenol-Free Neurobasal media (Gibco, 12348-017) supplemented with 2 mM GlutaMAX (Gibco 35050061) and B27 (Gibco 17504-044).

### Statistical analysis

One-way ANOVA with Tukey’s or Games-Howell’s post-hoc tests (depending on the condition of homoscedasticity and heteroscedasticity) was used for three or more dataset quantifications. Statistical calculations were performed with GraphPad Prism 7 Software (http://www.graphpad.com), *P* values <0.05 were considered significant. To correct for multiple comparisons in MS data analysis and neurite branching, the two-stage linear step-up procedure of Benjamini, Krieger and Yekutieli post-hoc tests was used to control the false discovery rate (FDR), q<0.05. Results are expressed as average + the standard error mean (SEM). A minimum of 3 independent biological replicates were used for each experiment with at least 3 replicates per sample within each experiment.

## ACKNOWLEDGEMENTS

We acknowledge Farida Korobova and Yvette Wong for critical reading of the manuscript.

## AUTHOR CONTRIBUTIONS

Conceptualization: GC

Methodology: EG, SZ, AG, GC

Investigation: EG, SZ, AG, GC

Visualization: EG, SZ

Supervision: GC

Writing—original draft: GC

Writing—review & editing: EG, GC

## DATA AND MATERIALS AVAILABILITY

All data are available in the main text or the supplementary materials.

## REFERENCES

1. A. Green, T. Hossain, D. M. Eckmann, Mitochondrial dynamics involves molecular and mechanical events in motility, fusion and fission. Front Cell Dev Biol 10, 1010232 (2022).

2. K. J. De Vos, V. J. Allan, A. J. Grierson, M. P. Sheetz, Mitochondrial function and actin regulate dynamin-related protein 1-dependent mitochondrial fission. Curr Biol 15, 678–683 (2005).

3. S. L. Archer, Mitochondrial fission and fusion in human diseases. N Engl J Med 370, 1074–1074 (2014).

4. Y. Yoon, K. R. Pitts, M. A. McNiven, Mammalian dynamin-like protein DLP1 tubulates membranes. MBoC 12, 2894–2905 (2001).

5. J. R. Friedman et al., ER tubules mark sites of mitochondrial division. Science 334, 358–362 (2011).

6. F. Korobova, V. Ramabhadran, H. N. Higgs, An actin-dependent step in mitochondrial fission mediated by the ER-associated formin INF2. Science 339, 464–467 (2013).

7. F. Korobova, T. J. Gauvin, H. N. Higgs, A role for myosin II in mammalian mitochondrial fission. Curr Biol 24, 409–414 (2014).

8. A. L. Hatch, W. K. Ji, R. A. Merrill, S. Strack, H. N. Higgs, Actin filaments as dynamic reservoirs for Drp1 recruitment. Mol Biol Cell 27, 3109–3121 (2016).

9. A. S. Moore, Y. C. Wong, C. L. Simpson, E. L. Holzbaur, Dynamic actin cycling through mitochondrial subpopulations locally regulates the fission-fusion balance within mitochondrial networks. Nat Commun 7, 12886 (2016).

10. S. Li et al., Transient assembly of F-actin on the outer mitochondrial membrane contributes to mitochondrial fission. J Cell Biol 208, 109–123 (2015).

11. U. Manor et al., A mitochondria-anchored isoform of the actin-nucleating spire protein regulates mitochondrial division. Elife 4, (2015).

12. W. K. Ji, A. L. Hatch, R. A. Merrill, S. Strack, H. N. Higgs, Actin filaments target the oligomeric maturation of the dynamin GTPase Drp1 to mitochondrial fission sites. Elife 4, e11553 (2015).

13. R. Chakrabarti et al., INF2-mediated actin polymerization at the ER stimulates mitochondrial calcium uptake, inner membrane constriction, and division. Journal of Cell Biology 217, 251–268 (2018).

14. A. S. Moore et al., Actin cables and comet tails organize mitochondrial networks in mitosis. Nature 591, 659–664 (2021).

15. T. S. Fung et al., Parallel kinase pathways stimulate actin polymerization at depolarized mitochondria. Curr Biol 32, 1577–1592. e1578 (2022).

16. S. M. Coscia et al., Myo19 tethers mitochondria to endoplasmic reticulum-associated actin to promote mitochondrial fission. J Cell Sci 136, (2023).

17. P. Gatti, C. Schiavon, J. Cicero, U. Manor, M. Germain, Mitochondria- and ER-associated actin are required for mitochondrial fusion. Nat Commun 16, 451 (2025).

18. D. M. Arduíno, A. R. Esteves, S. M. Cardoso, Mitochondrial fusion/fission, transport and autophagy in Parkinson′ s disease: When mitochondria get nasty. Parkinsons Dis 2011, 767230 (2011).

19. C. M. Deus et al., A mitochondria-targeted caffeic acid derivative reverts cellular and mitochondrial defects in human skin fibroblasts from male sporadic Parkinson’s disease patients. Redox Biol 45, 102037 (2021).

20. G. M. Lawrence, C. L. Holley, K. Schroder, Parkinson’s disease: connecting mitochondria to inflammasomes. Trends Immunol 43, 877–885 (2022).

21. F. Angelucci et al., Dementia with Lewy Bodies (DLB), Parkinson’s Disease (PD), and Multiple System Atrophy (MSA) Are Synucleopathies Characterized by Increased Serum Levels of Plasminogen Activator Inhibitor-1 (PAI-1). ACS omega, (2025).

22. B. M. Dufty et al., Calpain-cleavage of alpha-synuclein: connecting proteolytic processing to disease-linked aggregation. Am J Pathol 170, 1725–1738 (2007).

23. G. Caraveo et al., FKBP12 contributes to alpha-synuclein toxicity by regulating the calcineurin-dependent phosphoproteome. Proc Natl Acad Sci U S A 114, E11313–E11322 (2017).

24. G. Caraveo et al., Calcineurin determines toxic versus beneficial responses to alpha-synuclein. Proc Natl Acad Sci U S A 111, E3544–3552 (2014).

25. Z. S. Martin et al., alpha-Synuclein oligomers oppose long-term potentiation and impair memory through a calcineurin-dependent mechanism: relevance to human synucleopathic diseases. J Neurochem 120, 440–452 (2012).

26. L. F. Burbulla et al., Dopamine oxidation mediates mitochondrial and lysosomal dysfunction in Parkinson’s disease. Science 357, 1255–1261 (2017).

27. P. J. Teravskis et al., A53T Mutant Alpha-Synuclein Induces Tau-Dependent Postsynaptic Impairment Independently of Neurodegenerative Changes. J Neurosci 38, 9754–9767 (2018).

28. A. Shum et al., Octopamine metabolically reprograms astrocytes to confer neuroprotection against alpha-synuclein. Proc Natl Acad Sci U S A 120, e2217396120 (2023).

29. J. Ueda et al., Ca2+–Calmodulin–Calcineurin Signaling Modulates α-Synuclein Transmission. Movement Disorders 38, 1056–1067 (2023).

30. F. Widmer, P. Caroni, Identification, localization, and primary structure of CAP-23, a particle-bound cytosolic protein of early development. J Cell Biol 111, 3035–3047 (1990).

31. P. Caroni, L. Aigner, C. Schneider, Intrinsic neuronal determinants locally regulate extrasynaptic and synaptic growth at the adult neuromuscular junction. J Cell Biol 136, 679–692 (1997).

32. K. Mac Donald, A. Iulianella, The actin-cytoskeleton associating protein BASP1 regulates neural progenitor localization in the neural tube. genesis 60, e23464 (2022).

33. D. Frey, T. Laux, L. Xu, C. Schneider, P. Caroni, Shared and unique roles of CAP23 and GAP43 in actin regulation, neurite outgrowth, and anatomical plasticity. J Cell Biol 149, 1443–1454 (2000).

34. Y. Yamamoto, Y. Sokawa, S. Maekawa, Biochemical evidence for the presence of NAP-22, a novel acidic calmodulin binding protein, in the synaptic vesicles of rat brain. Neurosci Lett 224, 127–130 (1997).

35. S. Mason, Lactate Shuttles in Neuroenergetics-Homeostasis, Allostasis and Beyond. Front Neurosci 11, 43 (2017).

36. Q. Wang et al., The landscape of multiscale transcriptomic networks and key regulators in Parkinson’s disease. Nat Commun 10, 5234 (2019).

37. E. Grebenik, S. Zaichick, G. Caraveo, Calcineurin-mediated regulation of growth-associated protein 43 is essential for neurite and synapse formation and protects against α-synuclein-induced degeneration. Front Aging Neurosci 17, 1566465 (2025).

38. L. A. Greene, A. S. Tischler, Establishment of a noradrenergic clonal line of rat adrenal pheochromocytoma cells which respond to nerve growth factor. Proc Natl Acad Sci U S A 73, 2424–2428 (1976).

39. I. Korshunova et al., Characterization of BASP1-mediated neurite outgrowth. J Neurosci Res 86, 2201–2213 (2008).

40. T. Laux et al., GAP43, MARCKS, and CAP23 modulate PI (4, 5) P2 at plasmalemmal rafts, and regulate cell cortex actin dynamics through a common mechanism. The Journal of cell biology 149, 1455-1472 (2000).

41. M. M. Harraz et al., A high-affinity cocaine binding site associated with the brain acid soluble protein 1. Proc Natl Acad Sci U S A 119, e2200545119 (2022).

42. C. D. Nobes, A. Hall, Rho, rac, and cdc42 GTPases regulate the assembly of multimolecular focal complexes associated with actin stress fibers, lamellipodia, and filopodia. Cell 81, 53–62 (1995).

43. N. Martinez-Lopez et al., mTORC2-NDRG1-CDC42 axis couples fasting to mitochondrial fission. Nat Cell Biol 25, 989–1003 (2023).

44. E. Toska, J. Shandilya, S. J. Goodfellow, K. F. Medler, S. G. Roberts, Prohibitin is required for transcriptional repression by the WT1-BASP1 complex. Oncogene 33, 5100–5108 (2014).

45. K. Rajalingam et al., Prohibitin is required for Ras-induced Raf–MEK–ERK activation and epithelial cell migration. Nat Cell Biol 7, 837–843 (2005).

46. A. Signorile, G. Sgaramella, F. Bellomo, D. De Rasmo, Prohibitins: A Critical Role in Mitochondrial Functions and Implication in Diseases. Cells 8, (2019).

47. C. Merkwirth et al., Prohibitins control cell proliferation and apoptosis by regulating OPA1-dependent cristae morphogenesis in mitochondria. Genes Dev 22, 476–488 (2008).

48. K. Kasashima, E. Ohta, Y. Kagawa, H. Endo, Mitochondrial functions and estrogen receptor-dependent nuclear translocation of pleiotropic human prohibitin 2. J Biol Chem 281, 36401–36410 (2006).

49. K. H. Berger, M. P. Yaffe, Prohibitin family members interact genetically with mitochondrial inheritance components in Saccharomyces cerevisiae. Mol Cell Biol 18, 4043–4052 (1998).

50. C. Osman et al., The genetic interactome of prohibitins: coordinated control of cardiolipin and phosphatidylethanolamine by conserved regulators in mitochondria. J Cell Biol 184, 583–596 (2009).

51. R. Quintana-Cabrera, L. Scorrano, Determinants and outcomes of mitochondrial dynamics. Mol Cell 83, 857–876 (2023).

52. T. S. Fung, W. K. Ji, H. N. Higgs, R. Chakrabarti, Two distinct actin filament populations have effects on mitochondria, with differences in stimuli and assembly factors. J Cell Sci 132, (2019).

53. S. B. Song, J. S. Park, S. Y. Jang, E. S. Hwang, Nicotinamide Treatment Facilitates Mitochondrial Fission through Drp1 Activation Mediated by SIRT1-Induced Changes in Cellular Levels of cAMP and Ca2+. Cells 10, 612 (2021).

54. S. C. Leite, R. Pinto-Costa, M. M. Sousa, Actin dynamics in the growth cone: a key player in axon regeneration. Curr Opin Neurobiol 69, 11–18 (2021).

55. M. Schaks, G. Giannone, K. Rottner, Actin dynamics in cell migration. Essays Biochem 63, 483–495 (2019).

56. Z.-H. Sheng, Q. Cai, Mitochondrial transport in neurons: impact on synaptic homeostasis and neurodegeneration. Nat Rev Neurosci 13, 77–93 (2012).

57. H. Otera et al., Mff is an essential factor for mitochondrial recruitment of Drp1 during mitochondrial fission in mammalian cells. J Cell Biol 191, 1141–1158 (2010).

58. C. S. Palmer et al., Adaptor proteins MiD49 and MiD51 can act independently of Mff and Fis1 in Drp1 recruitment and are specific for mitochondrial fission. J Biol Chem 288, 27584–27593 (2013).

59. J. J. Collier, M. Oláhová, T. G. McWilliams, R. W. Taylor, Mitochondrial signalling and homeostasis: from cell biology to neurological disease. Trends Neurosci 46, 137–152 (2023).

60. J. Jackson et al., Actin-nucleation promoting factor N-WASP influences alpha-synuclein condensates and pathology. Cell Death Dis 15, 304 (2024).

61. D. G. Ordonez, M. K. Lee, M. B. Feany, α-synuclein induces mitochondrial dysfunction through spectrin and the actin cytoskeleton. Neuron 97, 108–124. e106 (2018).

62. J. Schindelin et al., Fiji: an open-source platform for biological-image analysis. Nat Methods 9, 676-682 (2012).

