## Supplemental Figures S1-S5 for "BASP1 Couples Ca^2+^ Signaling and Actin Polymerization to Mitochondrial Fission Essential for Neurite Outgrowth"

Ekaterina Grebenik *et al.*

**This PDF file includes:**

Figs. S1 to S5

**Other Supplementary Materials for this manuscript include the following:**

Movies S1 to S7
Table S1

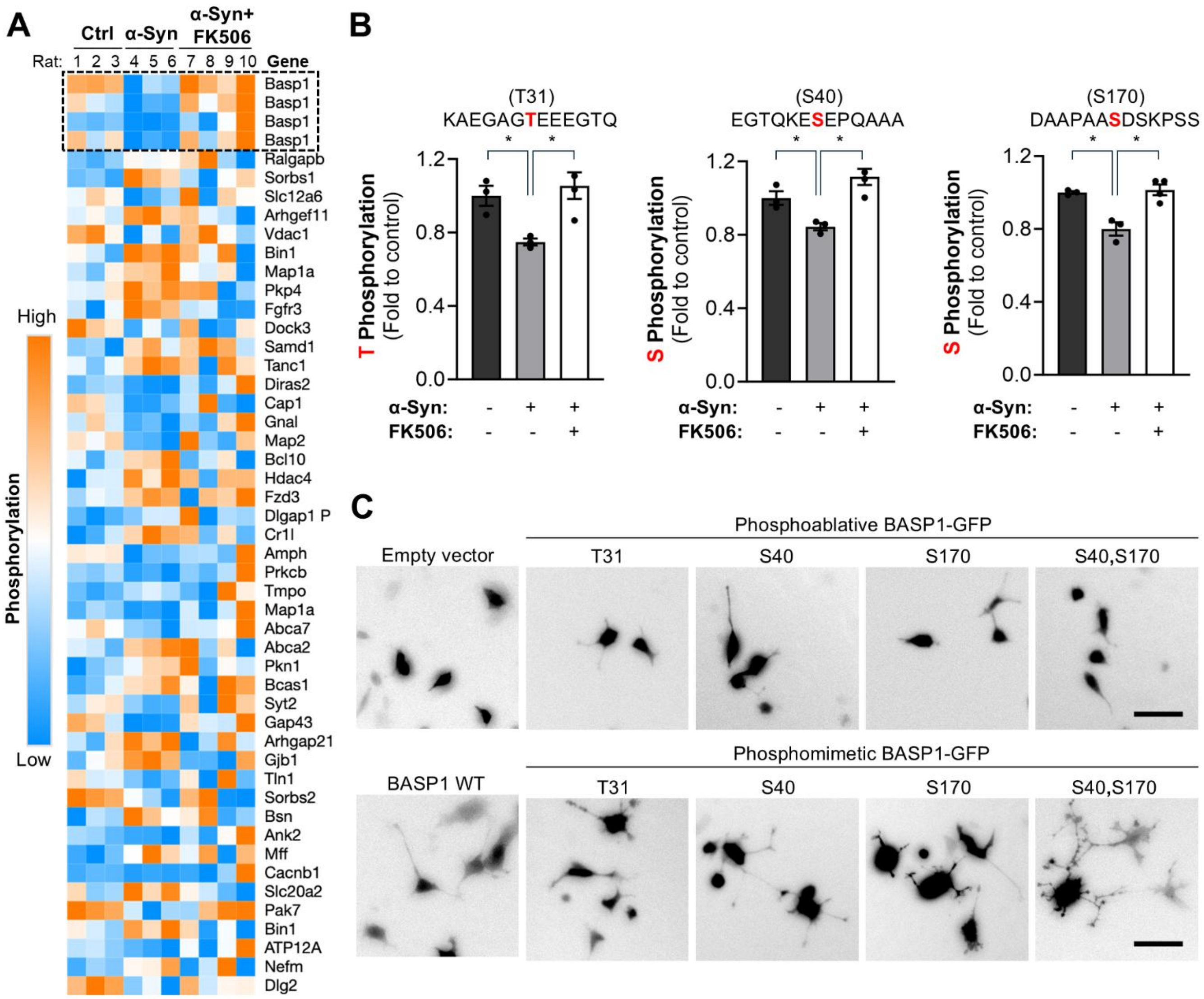
A
Ctrl
α-Syn
α-Syn+ FK506
Rat: 1 2 3 4 5 6 7 8 9 10
Gene
Basp1
Basp1
Basp1
Basp1
Ralgapb
Sorbs1
Slc12a6
Arhgef11
Vdac1
Bin1
Map1a
Pkp4
Fgfr3
Dock3
Samd1
Tanc1
Diras2
Cap1
Gnal
Map2
Bcl10
Hdac4
Fzd3
Dlgap1 P
Cr1l
Amph
Prkcb
Tmpo
Map1a
Abca7
Abca2
Pkn1
Bcas1
Syt2
Gap43
Arhgap21
Gjb1
Tln1
Sorbs2
Bsn
Ank2
Mff
Cacnb1
Slc20a2
Pak7
Bin1
ATP12A
Nefm
Dlg2
High
Phosphorylation
Low
B
(T31)
KAEGAGTEEEGTQ
T Phosphorylation (Fold to control)
(S40)
EGTQKESEPQAAA
S Phosphorylation (Fold to control)
(S170)
DAAPAASDSKPSS
S Phosphorylation (Fold to control)
α-Syn:
FK506:
C
Phosphoablative BASP1-GFP
Empty vector
T31
S40
S170
S40,S170
Phosphomimetic BASP1-GFP
BASP1 WT
T31
S40
S170
S40,S170

**Fig S1. Calcineurin-dependent phosphosites S40 and S170 of human BASP1 regulate neurite branching in PC12 cells. (A)** Heat map representing phosphoproteomic hits identified in the rat striatum after correction for protein abundance, showing statistically significant differences between control and α-synuclein (α-syn)-expressing rats *(23)*. Each row represents a protein from which the corresponding phosphorylated peptide was detected. Peptides outlined by the black dash line indicate BASP1 phosphorylation sites that restored phosphorylation to control levels in α-syn-expressing rats treated with sub-saturating doses of FK506. **(B)** Quantification of BASP1 phosphopeptide phosphorylation in control, α-syn, and α-syn + FK506-treated rats. The phosphorylated residue is indicated in red. N=3. Two-tailed t-test, *$P<0.05$. **(C)** Representative confocal images of PC12 cells co-transfected with GFP for neurite visualization and the following BASP1 constructs: wild-type (WT), phosphomimetic and phosphoablative T31, S40, S170, or S40/S170 mutants. Scale bar = 20 µm.

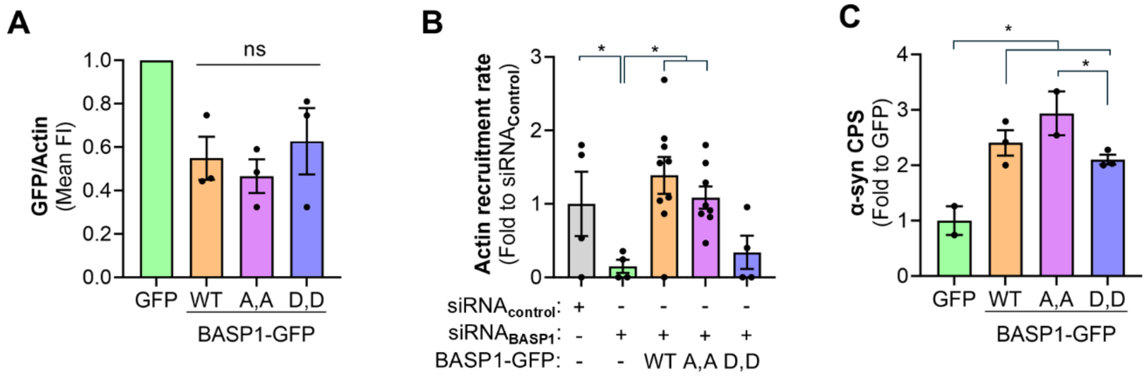
A
GFP/Actin (Mean FI)
ns
GFP WT A,A D,D
BASP1-GFP
B
Actin recruitment rate (Fold to siRNAControl)
siRNAcontrol:
siRNABASP1:
BASP1-GFP:
WT A,A D,D
C
α-syn CPS (Fold to GFP)
GFP WT A,A D,D
BASP1-GFP

**Fig. S2. Calcineurin-dependent phosphorylation sites S40 and S170 of BASP1 mediate binding to actin and α-synuclein. (A)** Quantitation of BASP1-GFP signal/actin, the loading control, from Fig. 1B, normalized to GFP alone within each experiment. N=3. One-way Anova with post hoc Tukey's test. **(B)** Quantification of the actin recruitment rate for Fig. 3K. One-way Anova with post hoc Dunnett's test, ****$P$<0.0001. **(C)** MS quantification of α-synuclein (α-syn) co-immunoprecipitated using anti-GFP beads from primary embryonic rat cortical neurons transduced with GFP alone or BASP1-GFP constructs: wild-type (WT), phosphoablative (S40A/S170A; A,A), and phosphomimetic (S40D/S170D; D,D) double mutants. One-Way Anova followed by multiple comparisons with two-stage linear step-up procedure of Benjamini, Krieger, and Yekutieli, *q<0.05

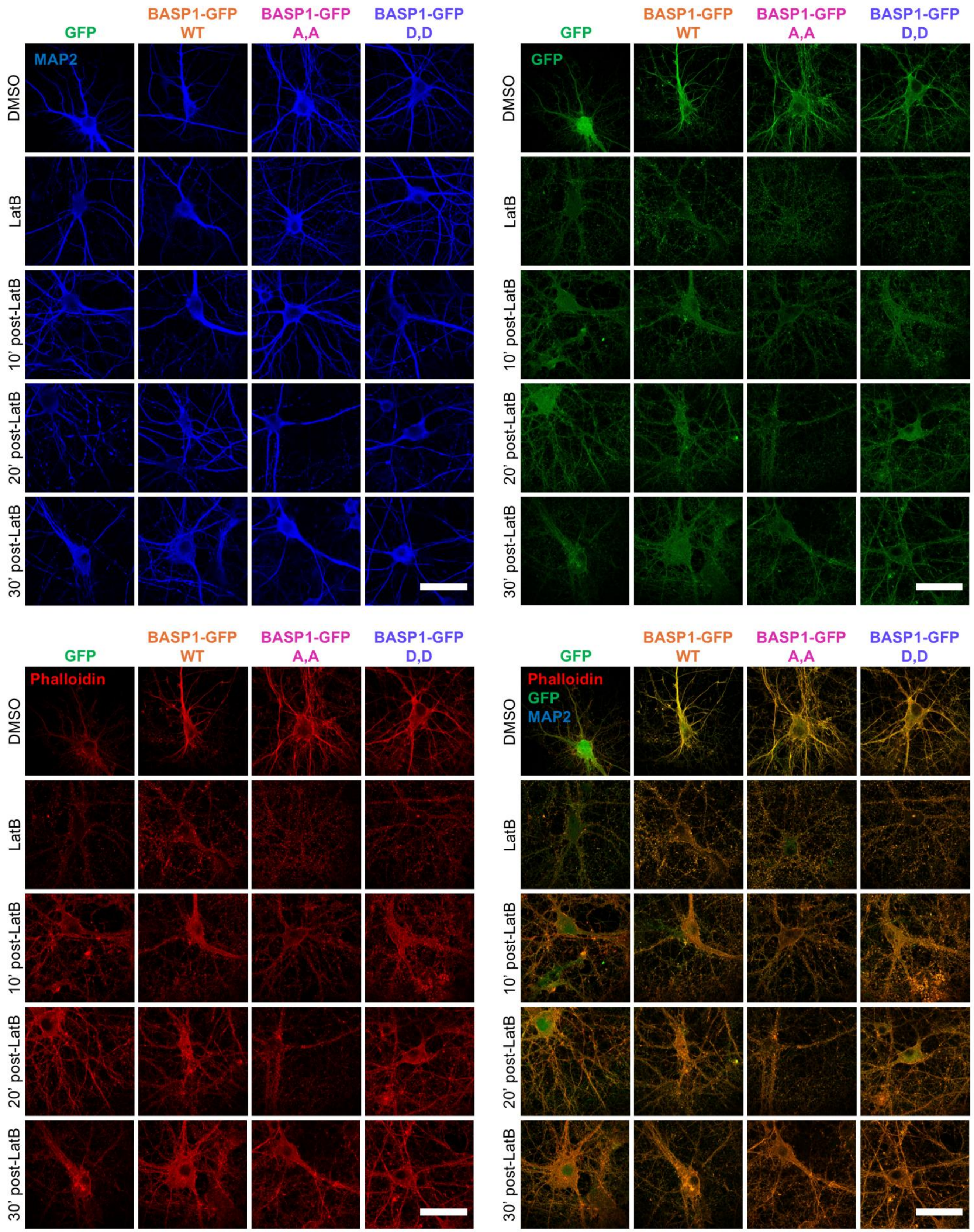
GFP
BASP1-GFP WT
BASP1-GFP A,A
BASP1-GFP D,D
MAP2
GFP
Phalloidin
Phalloidin
GFP
MAP2
DMSO
LatB
10' post-LatB
20' post-LatB
30' post-LatB

**Fig. S3. Calcineurin-dependent phosphorylation sites S40 and S170 of human BASP1 regulate actin polymerization in primary embryonic rat cortical neurons.** Representative confocal images of time course of actin polymerization following 10-minute 2 μM Latrunculin B (LatB) treatment of DIV23 primary embryonic rat cortical neurons transduced with GFP alone or BASP1-GFP constructs: wild-type (WT), phosphoablative (S40A/S170A; A,A), and phosphomimetic (S40D/S170D; D,D) double mutants. Neurons were identified by MAP2 immunostaining. F-actin was visualized using BODIPY™ 558/568 Phalloidin. Scale bar = 50 μm.

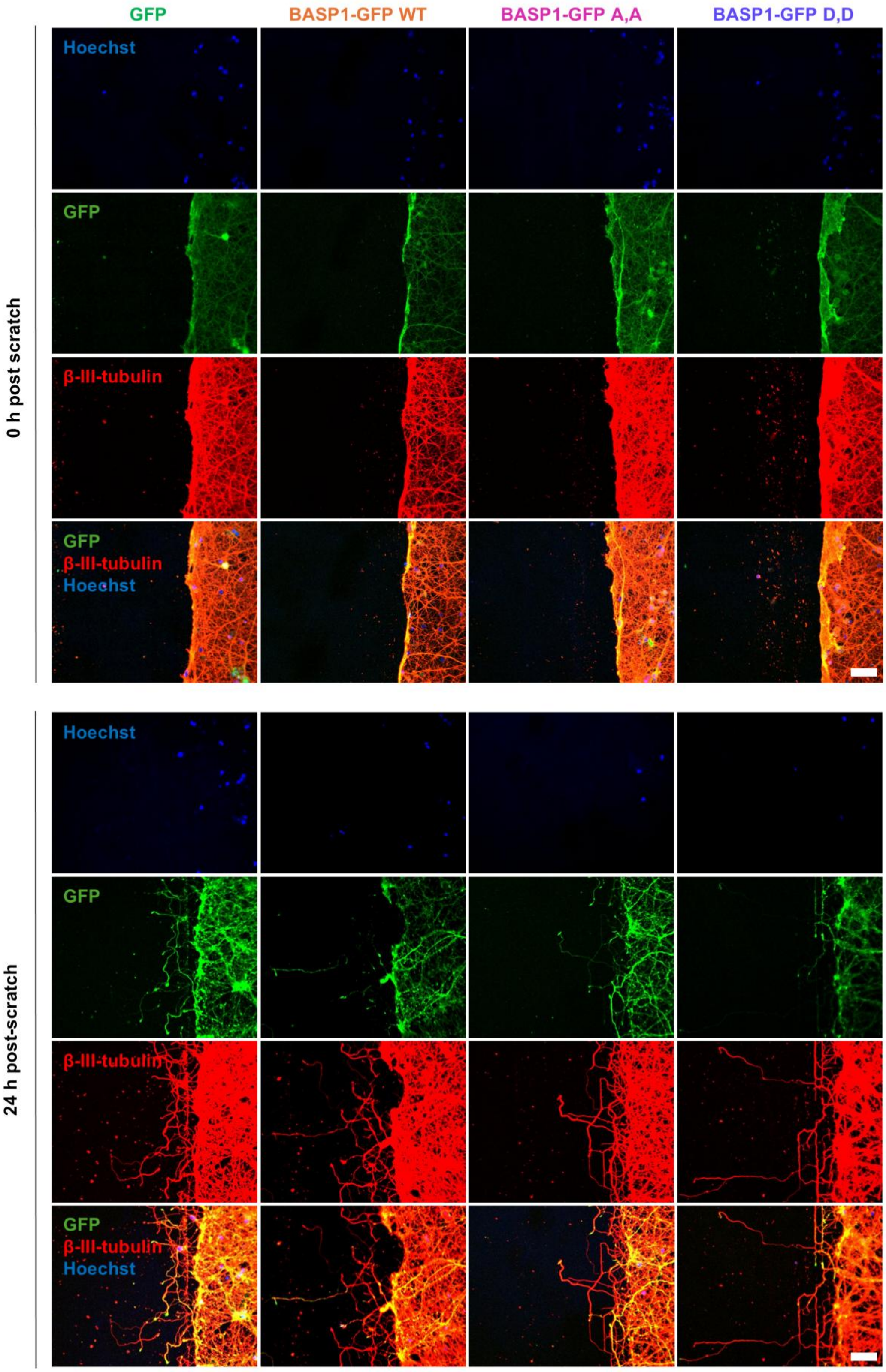

GFP
BASP1-GFP WT
BASP1-GFP A,A
BASP1-GFP D,D
0 h post scratch
Hoechst
GFP
β-III-tubulin
GFP
β-III-tubulin
Hoechst
24 h post-scratch
Hoechst
GFP
β-III-tubulin
GFP
β-III-tubulin
Hoechst

**Fig. S4. Calcineurin-dependent phosphorylation sites S40 and S170 of human BASP1 regulate neurite regeneration following mechanical injury in primary embryonic rat cortical neurons.** Representative confocal images of β-III-tubulin staining in DIV23 primary embryonic rat cortical neurons, transduced with either GFP alone or BASP1-GFP constructs: wild-type (WT), phosphoablative (S40A/S170A; A,A), and phosphomimetic (S40D/S170D; D,D) double mutants. Neurons were subjected to mechanical injury in a scratch assay, and the staining was performed 24 hours post-injury. Scale bar = 50 µm.

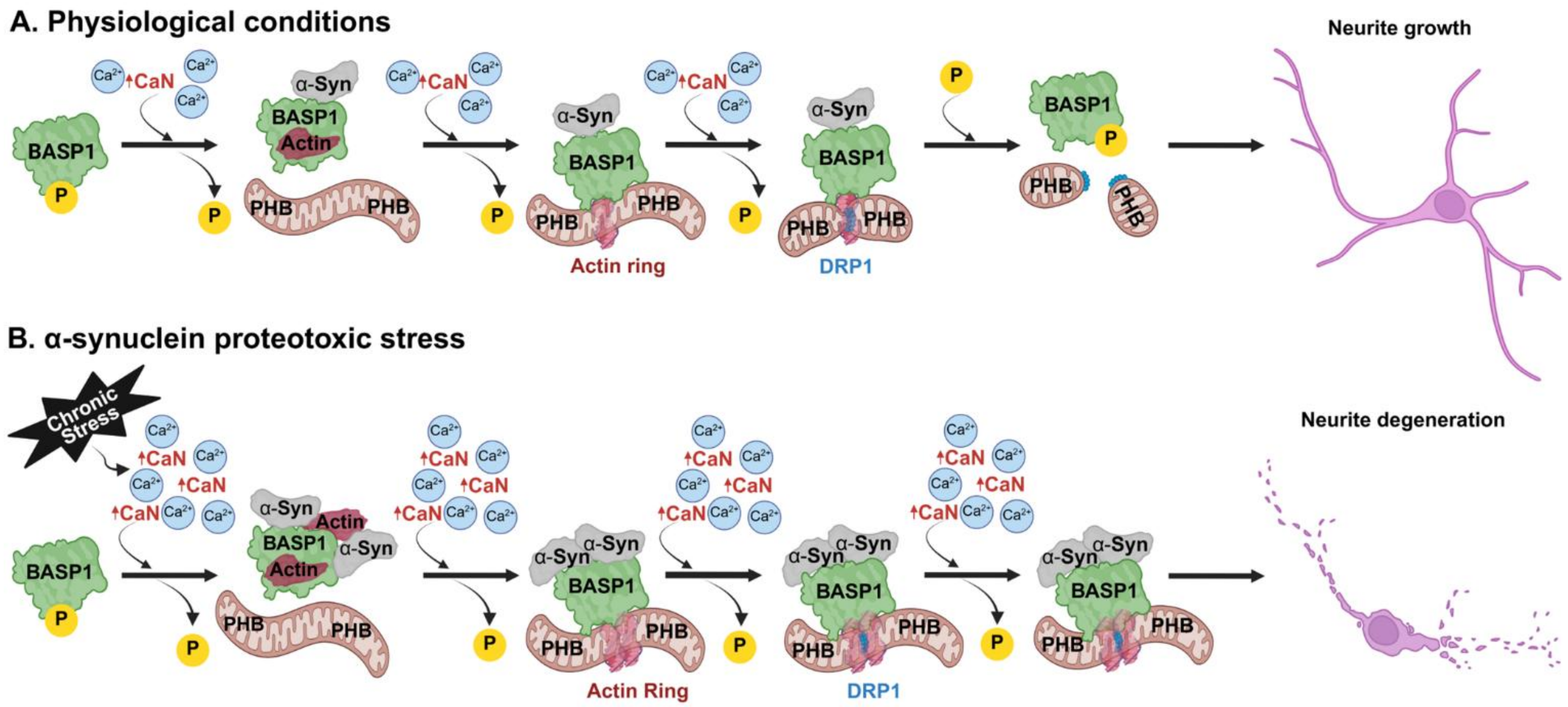

A. Physiological conditions
BASP1
P
Ca2+
CaN
α-Syn
Actin
PHB
Actin ring
DRP1
Neurite growth
B. α-synuclein proteotoxic stress
Chronic Stress
Actin Ring
Neurite degeneration

**Fig. S5. Graphic model of the BASP1-mediated regulation of α-synuclein (α-syn) pathology in $Ca^{2+}$-CaN-dependent manner.** Biorender software was used to create the figure under an academic license.

**Table S1.**

Phosphoproteomic profiling of the striatum by TMT-MS in a rat model of α-synucleinopathy treated with FK506.

**Movie S1.**

Representative time-lapse confocal microscopy of actin, wild-type (WT) BASP1, and mitochondria in HeLa cells co-transfected with BASP1-GFP WT and a 3′UTR-targeting siRNA against human BASP1. Cells were treated with 4 μM ionomycin. Actin was visualized via transient expression of LifeAct-mScarlet, and mitochondria were labeled with MitoTracker Deep Red. Images were acquired at 15-second intervals.

**Movie S2.**

Representative time-lapse confocal microscopy of actin, phosphoablative BASP1, and mitochondria in HeLa cells co-transfected with BASP1-GFP S40A/S170A double mutant and a 3′UTR-targeting siRNA against human BASP1. Cells were treated with 4 μM ionomycin. Actin was visualized via transient expression of LifeAct-mScarlet, and mitochondria were labeled with MitoTracker Deep Red. Images were acquired at 15-second intervals.

**Movie S3.**

Representative time-lapse confocal microscopy of actin, phosphomimetic BASP1, and mitochondria in HeLa cells co-transfected with BASP1-GFP S40D/S170D double mutant and a 3′UTR-targeting siRNA against human BASP1. Cells were treated with 4 μM ionomycin. Actin was visualized via transient expression of LifeAct-mScarlet, and mitochondria were labeled with MitoTracker Deep Red. Images were acquired at 15-second intervals.

**Movie S4.**

Representative time-lapse confocal microscopy of Drp1, wild-type (WT) BASP1, and mitochondria in HeLa cells co-transfected with BASP1-GFP WT and a 3′UTR-targeting siRNA against human BASP1. Cells were treated with 4 μM ionomycin. Drp1 was visualized via transient expression of Drp1-mCherry, and mitochondria were labeled with MitoTracker Deep Red. Images were acquired at 15-second intervals.

**Movie S5.**

Representative time-lapse confocal microscopy of Drp1, phosphomimetic BASP1, and mitochondria in HeLa cells co-transfected with BASP1-GFP S40D/S170D double mutant and a 3′UTR-targeting siRNA against human BASP1. Cells were treated with 4 μM ionomycin. Drp1 was visualized via transient expression of Drp1-mCherry, and mitochondria were labeled with MitoTracker Deep Red. Images were acquired at 15-second intervals.

**Movie S6.**

Representative time-lapse confocal microscopy of Drp1, phosphoablative BASP1, and mitochondria in HeLa cells co-transfected with BASP1-GFP S40A/S170A double mutant and a 3′UTR-targeting siRNA against human BASP1. Cells were treated with 4 μM ionomycin. Drp1 was visualized via transient expression of Drp1-mCherry, and mitochondria were labeled with MitoTracker Deep Red. Images were acquired at 15-second intervals.

**Movie S7.**

Representative time-lapse confocal microscopy of BASP1 and mitochondria in primary embryonic rat cortical neurons transduced with BASP1-GFP constructs: wild-type (WT), phosphoablative (S40A/S170A; A,A), and phosphomimetic (S40D/S170D; D,D) double mutants. Mitochondria were visualized in neurites with the fluorescent mitochondrial dye tetramethylrhodamine methyl ester (TMRM) 24 hours post-injury. Images were acquired at 3-min intervals.
